# A tripartite protein complex promotes DNA transport during natural transformation in firmicutes

**DOI:** 10.1101/2025.05.09.652193

**Authors:** Marie Dewailly, Yoann Fauconnet, Cécile Ducrot, Anne-Lise Soulet, Nathalie Campo, Raphael Guerois, J. Pablo Radicella, Patrice Polard, Jessica Andreani, Calum Johnston

## Abstract

Natural genetic transformation is a conserved mechanism of bacterial horizontal gene transfer, which is directed entirely by the recipient cell and facilitates the acquisition of new genetic traits such as antibiotic resistance. Transformation proceeds via the capture of exogenous DNA, its internalisation in single strand form (ssDNA) and its integration into the recipient chromosome by homologous recombination. While the proteins involved in these steps have mainly been identified, the specific mechanisms at play remain poorly characterised. This study takes advantage of recent advances in structural modelling to explore the uptake of ssDNA during transformation. Using the monoderm human pathogen *Streptococcus pneumoniae*, we model a tripartite protein complex composed of the transmembrane channel ComEC, and two cytoplasmic ssDNA-binding proteins ComFA and ComFC. Using targeted mutation and transformation assays, we propose that pneumococcal ComEC features a narrow channel for ssDNA passage, and we show this channel is conserved in the diderm *Helicobacter pylori*. We identify key residues involved in protein-protein and protein-ssDNA interactions in the pneumococcal tripartite complex model and we show them to be crucial for transformation efficiency. Structural modelling reveals that this tripartite protein complex and its interaction with ssDNA are conserved in firmicutes. Overall, this study validates a tripartite complex required for the internalisation of ssDNA during transformation in firmicutes, providing new insights into the molecular mechanisms involved in this horizontal gene transfer mechanism central to bacterial adaptation. It also demonstrates the power of recent structural modelling techniques such as AlphaFold3 as hypothesis generators and guides for designing experiments.

**Significance statement:** Natural genetic transformation is a key mechanism of horizontal gene transfer, conserved in bacteria. The investigation of transmembrane channel ComEC and its interaction partners was previously hindered by the difficulty of manipulating these proteins experimentally. Thanks to state-of-the-art structural modelling with AlphaFold3 and subsequent thorough experimental validation of this model using targeted mutation and transformation assays, we demonstrate the importance of the ComEC/ComFA/ComFC complex for single strand DNA (ssDNA) uptake during natural transformation in the human pathogen *Streptococcus pneumoniae*. Similar models in several other species suggest a widely conserved organization of this complex in firmicutes. In addition, we demonstrate that the ComEC transmembrane channel is also crucial for ssDNA uptake during natural transformation in *Helicobacter pylori*.

## Introduction

Natural genetic transformation is a mechanism of horizontal gene transfer (HGT) well conserved in the bacterial kingdom^1^. Transformation involves the capture of exogenous DNA, followed by its internalisation in single strand form (ssDNA) and its integration into the bacterial chromosome by homologous recombination. Transformation is the only HGT mechanism directed entirely by the recipient cell, and promotes the acquisition of new genetic traits, facilitating the spread of antibiotic resistance^2^. Transformation occurs during a physiological state called competence^1,3,4^, where the genes encoding transformation proteins are induced^5–7^. Although the regulation of competence varies widely across bacterial species, the key steps of transformation and some of the central proteins involved are broadly conserved^1^. These steps include DNA capture, facilitated in most cases by a transformation pilus^8–13^, ssDNA import across the cytoplasmic membrane mediated by the ComEC membrane channel^1,14^, and chromosomal integration by homologous recombination, ensured by the conserved homologous recombinase RecA and its transformation-dedicated recombination mediator protein DprA^15–17^. These three steps are universal to all transformable species.

Although the key proteins involved in transformation have largely been identified, many gaps in the understanding of the molecular mechanisms and interactions involved in transformation remain, including at the stage of ssDNA import across the cytoplasmic membrane. The main actor in this step is the ComEC transmembrane protein, which is thought to form a channel for ssDNA to pass through and as such is essential for DNA uptake during transformation^14,18,19^. The ComEC protein, characterised by the universal presence of the *Competence* domain^20^, is conserved in all transformable species^1^, and thought to be ubiquitous in bacteria. Indeed, although the transmembrane *Competence* domain is crucial for transformation, it is also present in many bacterial species not shown to be transformable. In addition, two other domains, an OB-fold and a β–lactamase domain, are variably present in the ComEC protein, with different configurations present across transformable species^20^. While the transmembrane *Competence* domain is thought to form the channel through which transforming ssDNA passes to cross the bacterial membrane, the roles of the two other domains in transformation remain unclear. In the firmicute model organism *Bacillus subtilis*, the ComEC protein (*Bs*ComEC) possesses both an OB-fold and a β–lactamase domain, with the latter proposed to participate in the degradation of one strand of transforming DNA^21,22^. This is also the case in the Gram-negative Gammaproteobacteria species *Legionella pneumophila*^23^. However, this is not the case in the pathogenic firmicute model *Streptococcus pneumoniae*, where this role is carried out by the EndA nuclease^24–26^, despite the pneumococcal ComEC protein (*Sp*ComEC) possessing a (possibly inactive) β–lactamase domain as well as an OB-fold^1,20,21^. In addition, since neither the OB-fold or a β–lactamase domain are conserved across transformable species, these domains cannot play universally conserved roles in transformation.

Another protein universal in transformable species is ComFC (also called ComF in some species), which is an ssDNA-binding protein key for transformation and shown to be involved in ssDNA import in the diderm pathogen *Helicobacter pylori*^27^, but whose specific role in transformation has remained elusive. In firmicutes, a third protein is proposed to be involved in ssDNA uptake, the ATP-dependent ssDNA helicase ComFA, which is suggested to pull transforming ssDNA through the *Sp*ComEC channel^28,29^. Pneumococcal ComFA (*Sp*ComFA) and ComFC (*Sp*ComFC) have been shown to interact in bacterial two-hybrid assays^28^. Like *Sp*ComEC, *Sp*ComFA was shown to be necessary for ssDNA uptake during pneumococcal transformation^19^. Together, these studies place *Sp*ComFA and *Sp*ComFC at the interface between ssDNA internalisation and subsequent chromosomal integration in firmicutes, and suggest that the *Sp*ComFA/FC protein pair may interact with *Sp*ComEC to ensure the import of ssDNA during transformation (Figure 1A). One hindrance to our understanding of ssDNA import during transformation is the difficulty in experimental manipulation of the ComEC transformation channel protein, which is predicted to be embedded in the cytoplasmic membrane^14^. Due to its hydrophobic nature, its size and its membrane association, biochemical and structural studies of ComEC are complex. In addition, *spcomEC* is expressed at relatively low levels during competence, and the *Sp*ComEC protein is rapidly degraded by the membrane-associated serine protease HtrA^30^.

**Figure 1:**
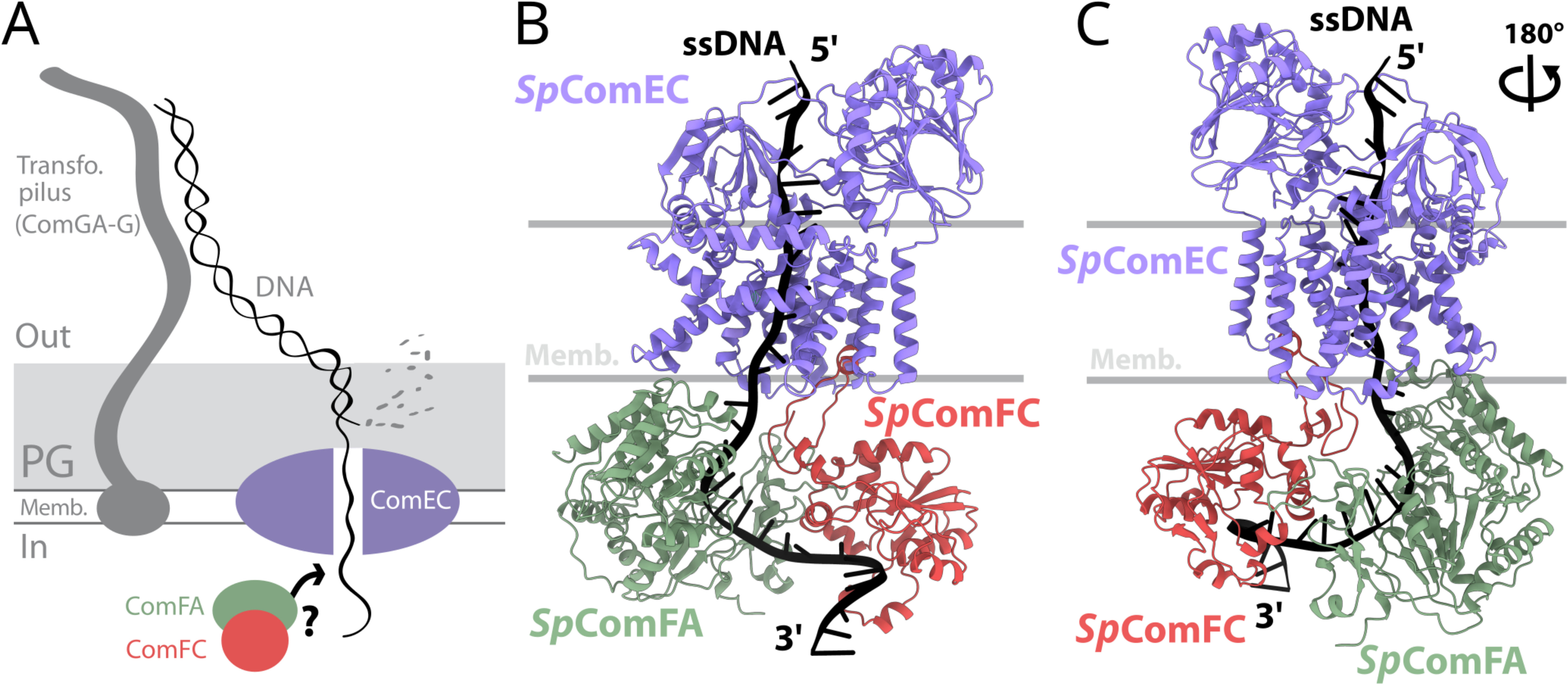
A model of the tripartite complex transporting ssDNA across the membrane during pneumococcal transformation. (A) DNA uptake machinery of the pneumococcal transformasome. Transforming DNA initially interacts with a transformation pilus made up of the ComGA-G proteins^8–10^. Retraction of this pilus passes the DNA to the DNA receptor ComEA^19,64^, and the EndA nuclease degrades one strand of the DNA^19,25^, with the remaining intact ssDNA strand passing through *Sp*ComEC into the cytoplasm. Highlighted in colour are *Sp*ComEC, a membrane protein proposed to provide an ssDNA channel, as well as *Sp*ComFA and *Sp*ComFC, cytoplasmic proteins known to form a complex and thought to be involved in this process. (B) Structural model of a tripartite complex between pneumococcal *Sp*ComEC (UniprotID: Q8DQ40, in purple), *Sp*ComFA (UniprotID: Q8CWM9, in green), *Sp*ComFC (UniprotID: Q8CWN0, in red) and ssDNA (30-bp poly-A ssDNA, in black, with polarity shown). Model generated using AlphaFold3^2^, Model confidence: ipTM, 0.74; pTM, 0.82. These values evaluate the confidence of the predicted protein structure and complexes. Inter-chain predicted Template Modelling score (ipTM) reflects confidence in multiprotein predictions (0-1 range, with higher values representing higher confidence in the interaction interfaces). Predicted Template Modelling score (pTM) is a confidence metric predicting how the overall predicted fold will reflect true structure (0-1 range, with higher values representing confidence in predicted protein structure). (C) Tripartite complex structural model from panel B rotated 180° on the vertical axis.

In this study, we addressed this challenging question by using a structural modelling approach via AlphaFold3^31^ to explore the predicted structure of the ComEC transmembrane channel and its cytoplasmic transformation partners. Using pneumococcal proteins, we identified a tripartite protein complex of *Sp*ComEC, *Sp*ComFA and *Sp*ComFC in interaction with ssDNA. This model predicted with high confidence that these three proteins interact with 1:1:1 stoichiometry, with *Sp*ComEC featuring a narrow channel allowing ssDNA to pass through the cell membrane. The organisation of the ComEC *Competence* domain as a monomeric transmembrane channel was found to be conserved across transformable species, as key residues are conserved in both monoderm (*S. pneumoniae*) and diderm (*H. pylori*) species. Similar tripartite protein models of ComEC-FA-FC interacting with ssDNA were observed for nine other firmicute species. Key residues involved in protein-protein and protein-ssDNA interactions were identified in the pneumococcal model, and validated using transformation assays. In particular, an interaction between ComEC and ComFC was found to be crucial for transformation, and conserved across firmicutes. These results reveal for the first time the molecular choreography involved in ssDNA import across the cytoplasmic membrane during transformation in bacteria, and we propose that this tripartite complex represents a conserved mode of ssDNA import for transformation in firmicutes.

## Results

### Structural model of a tripartite protein complex proposed to be involved in ssDNA import during pneumococcal transformation

To explore potential interactions between ComEC, ComFA and ComFC in firmicutes, a structural interaction model was generated with AlphaFold3^31^ using the human pathogen *S. pneumoniae* as a model organism. Using *Sp*ComEC, *Sp*ComFA and *Sp*ComFC as well as a 30-nt poly-A DNA molecule, we generated a high-confidence model (Figure 1BC) (ipTM = 0.74, pTM = 0.82). This model revealed a three-way interaction between *Sp*ComEC, *Sp*ComFA and *Sp*ComFC in complex, with ssDNA traversing a narrow channel within a *Sp*ComEC monomer, before interacting in turn with *Sp*ComFA and *Sp*ComFC.

Although the confidence in the ssDNA structure and its interactions in this model was relatively low (chain pTM: 0.7-0.88 for protein chains and 0.08 for ssDNA, ipTM for pairs of chains: 0.65-0.84 for protein-protein and 0.27-0.45 for protein-ssDNA interactions), the complementary electrostatic surfaces and evolutionary conservation of residues on the proteins at ssDNA interaction sites strongly support the relevance of the model (Figure S1). While ComEC and ComFC are conserved across all transformable species, the ComFA protein is conserved only in firmicutes (Table S1). Similar tripartite models from nine other firmicute species were predicted with a high confidence (Figure S2, ipTM from 0.71 to 0.76, pTM from 0.77 to 0.84) and showed remarkable robustness (Table S2), with RMSD ranging from 1.5 Å to 3.0 Å compared to *S. pneumoniae*, despite sequence identity as low as 20%, 30% and 32% for ComEC, ComFC and ComFA, respectively. These findings suggest that this complex, proposed to ensure ssDNA import during transformation, is conserved across firmicutes. The robustness of our modelling was further confirmed by a similar high confidence tripartite model, obtained for ComEC, ComFC and ComFA from *Bacillus subtilis* in a recent high-throughput study using AlphaFold2^32^. Unlike the present study, AlphaFold2 did not enable the modelling of ssDNA, and the *B. subtilis* ComEC-ComFC-ComFA model was not explored further.

### ComEC is a conserved monomeric transmembrane channel crucial for ssDNA import during transformation in monoderm and diderm bacteria

In the pneumococcus, the *spcomEC* gene is induced solely during competence^5–7^. To detect *Sp*ComEC via Western blot, an alfa tag^33^ was inserted into a cytoplasmic loop of *Sp*ComEC predicted via AlphaFold3 to have minimal impact on protein structure. This mutant (*Sp*ComEC^alfa349^) was found to be functional for transformation (Figure S3AB). Time-course Western blots using anti-alfa antibodies showed that *Sp*ComEC was expressed at maximal levels 8-10 minutes after competence induction (Figure S3CD). Focus on *Sp*ComEC in interaction with ssDNA in the structural model presented in Figure 1BC identified one transmembrane domain, discontinuous in sequence, as well as conserved OB-fold (CATH topology 2.40.50) and β-lactamase domains (CATH superfamily 3.60.15.10) (Figure 2A), both of which were positioned on the outside of the cell membrane (Figure 2B). ssDNA was predicted to pass through a narrow channel in an *Sp*ComEC monomer (Figure 2A). In all models of the tripartite complex with ssDNA, in *S. pneumoniae* (Figure 1) and in other firmicute species (Figure S2), ssDNA consistently passed through this channel in the same orientation, with the 3’ end entering the cell first, as previously proposed^26,34^.

**Figure 2:**
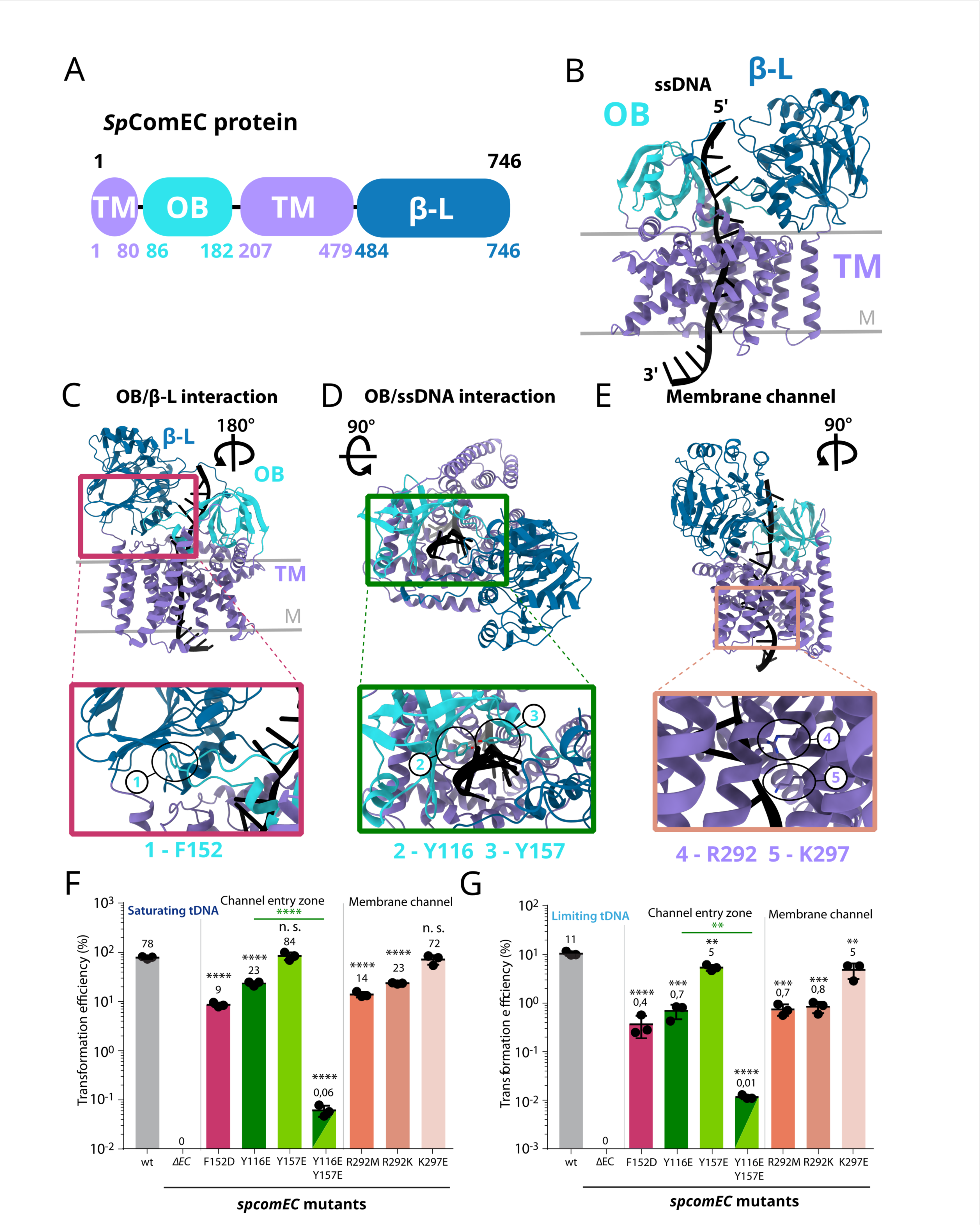
Mutation of selected residues on pneumococcal *Sp*ComEC protein affects transformation efficiency. (A) Domain organisation of *Sp*ComEC protein showing delimitations between two transmembrane domains (TM), an OB-fold (OB) and a β-lactamase domain (β-L). Numbers shown represent amino acids (AA) along the 746 AA protein. (B) Structural model of *Sp*ComEC interacting with ssDNA, with OB fold (light blue), β-lactamase domain (dark blue) and transmembrane domains (purple) shown. Grey lines represent the approximate position of the membrane. Polarity of ssDNA in the model is shown. (C) Highlight of F152 (1) on OB fold, predicted to interact with β-lactamase domain near the entry point of the membrane channel. (D) Highlight of Y116 (2) and Y157 (3) on OB fold, predicted to interact with transforming ssDNA near the entry point of the membrane channel. (E) Highlight of R292 (4) and K297 (5) in the transmembrane channel, predicted to interact with transforming ssDNA. (F) Transformation efficiency of *Sp*ComEC mutants when transformed with saturating concentrations of tDNA (2,500 pg mL^-1^ of a 3,434 bp PCR fragment possessing *rpsL41* point mutation, conferring streptomycin resistance^35,36^). Strains used: *wt*, R1501; *ΔSpcomEC*, R2300; F152D, R5003; Y116E, R5304; Y157E, R5001; Y116E-Y157E, R5002; R292M, R5007; R292K, R5009; K297E, R5011. Error bars representative of triplicate repeats, with means and individual data points shown. Student’s t-tests were used to compare transformation efficiencies of mutant strains with wildtype, unless shown by horizontal line. ****, p < 0.0001; n. s., not significant. (G) Transformation efficiency of *Sp*ComEC mutants when transformed with limiting concentrations of tDNA (25 pg mL^-1^). Strains used, tDNA identity, data representation and statistical analyses as in panel F. ****, p < 0.0001; ***, p < 0.001; **, p < 0.01.

To explore which interactions were important for the role of *Sp*ComEC in transformation, we identified residues predicted to be located at the interface between domains or interacting with ssDNA. Residue positions to be mutated were chosen based on their evolutionary conservation (Figure S3E), predicted involvement in the interactions between *Sp*ComEC domains or between *Sp*ComEC and ssDNA, and surface exposure, to minimize potentially disruptive effects on protein structure and stability. First, F152 was identified on the OB-fold and predicted to interact with the β-lactamase domain (Figure 2C). Second, Y116 and Y157, also present in the OB-fold, were predicted to interact with transforming ssDNA (Figure 2D). Finally, R292 and K297 were identified in the structural model as protruding into the membrane channel and interacting with transforming ssDNA (Figure 2E).

To test the functional importance of these residues, mutations predicted to disrupt these interactions were generated in a wildtype pneumococcal strain and compared in transformation assays. tDNA was a 3,434 bp PCR fragment possessing a centrally-located point mutation (*rpsL41*) conferring streptomycin resistance^35,36^. The tested mutants did not notably affect the stability of a *Sp*ComEC^alfa349^ fusion (Figure S3F). Under saturating tDNA conditions, transformation in the wildtype strain was 78 %, while a *ΔspcomEC* strain did not transform, confirming the essentiality of *Sp*ComEC for transformation. A *Sp*ComEC^F152D^ mutant showed a significant ∼9-fold decrease in transformation efficiency (Figure 2F). This suggests that the interaction between the OB-fold and β-lactamase domains is important for transformation. Then, *Sp*ComEC^Y116E^ and *Sp*ComEC^Y157E^ mutants were generated. While *Sp*ComEC^Y116E^ reduced transformation ∼4-fold, no effect was observed for *Sp*ComEC^Y157E^ (Figure 2F). However, a strain possessing both mutations displayed a dramatic 1,200-fold reduction in transformation, far larger than *Sp*ComEC^Y116E^ alone, suggesting that both residues contribute to the interaction between the OB-fold and transforming ssDNA. Finally, a series of mutants aiming to disrupt the passage of ssDNA through the membrane channel of *Sp*ComEC was generated. Mutation of the R292 position to either methionine (*Sp*ComEC^R292M^) or lysine (*Sp*ComEC^R292K^) resulted in a similar 5-10-fold decrease in transformation (Figure 2F). However, no effect was observed for a *Sp*ComEC^K297E^ mutant.

Under saturating tDNA conditions, cells are transformed with several tDNA molecules, each conferring resistance (Figure S4A), which may mask subtle effects of mutations on transformation efficiency. To address this, the experiments were repeated with limiting concentrations of tDNA (100-fold lower than in saturating conditions, resulting in decreased transformation efficiency in wildtype) (Figure S4B). Under these conditions, the effects of mutations were more pronounced, with significant differences even for the *Sp*ComEC^Y157E^ and *Sp*ComEC^K297E^ mutants (Figure 2G). Thus, conditions where tDNA molecules are less abundant reveal the importance of these residues in transformation. Taken together, these findings identify residues in *Sp*ComEC that are important for transformation. In addition, the fact that altering residues in the membrane channel affects transformation efficiency provides evidence that ssDNA passes through a *Sp*ComEC monomer into cells during transformation.

The ComEC transformation channel protein is conserved across all transformable species, although the OB-fold and β–lactamase domains are not universally present^20^. To determine if the monomeric membrane channel structure is conserved in diderm species, we generated an Alphafold3 model of ComEC from *H. pylori* (*Hp*ComEC) with ssDNA. The *Hp*ComEC model possesses a transmembrane domain, discontinuous in sequence, and an OB-fold, but no β-lactamase domain (Figure 3A). The AlphaFold3 structural model of *Hp*ComEC with a 15-nt poly-A DNA molecule, combined with evolutionary conservation information, enabled us to single out three residues predicted to interact with ssDNA: two residues in the OB-fold domain (Y60 and Y82) and one residue in the membrane channel (R239). Y82 and R239 in *Hp*ComEC are the equivalent of Y116 and R292 in *Sp*ComEC, respectively (Figures 2 and S5A). In contrast, the Y60 residue of *Hp*ComEC has no equivalent on *Sp*ComEC, but was chosen based on evolutionary conservation in *H. pylori* and related species (Figure S5B). In addition, the 3’-to-5’ polarity of ssDNA in the modelled *Hp*ComEC channel mirrored that observed in firmicutes. Transformation assays revealed that a *Hp*ComEC-FLAG mutant displayed wildtype transformation efficiency, while *Hp*ComEC^Y60E-Y82E^-FLAG, *Hp*ComEC^R239M^-FLAG and *Hp*ComEC^R239K^-FLAG mutants displayed significant deficits in transformation efficiency (Figure 3C), but remained above that of a Δ*hpcomEC* mutant^27^. Although *Hp*ComEC-FLAG proteins could not be detected by anti-FLAG Western blot at native levels, ectopic over-expression of these mutants in a *P_ureA_-HpcomEC-alfa* genetic context allowed their stability to be validated using anti-alfa antibodies (Figure S5C). These results confirm that these conserved residues are key for transformation in both *S. pneumoniae* and *H. pylori*. Taken together, these data support the model of ComEC as a conserved monomeric transmembrane channel ensuring the import of ssDNA during transformation in both monoderm and diderm bacteria.

**Figure 3:**
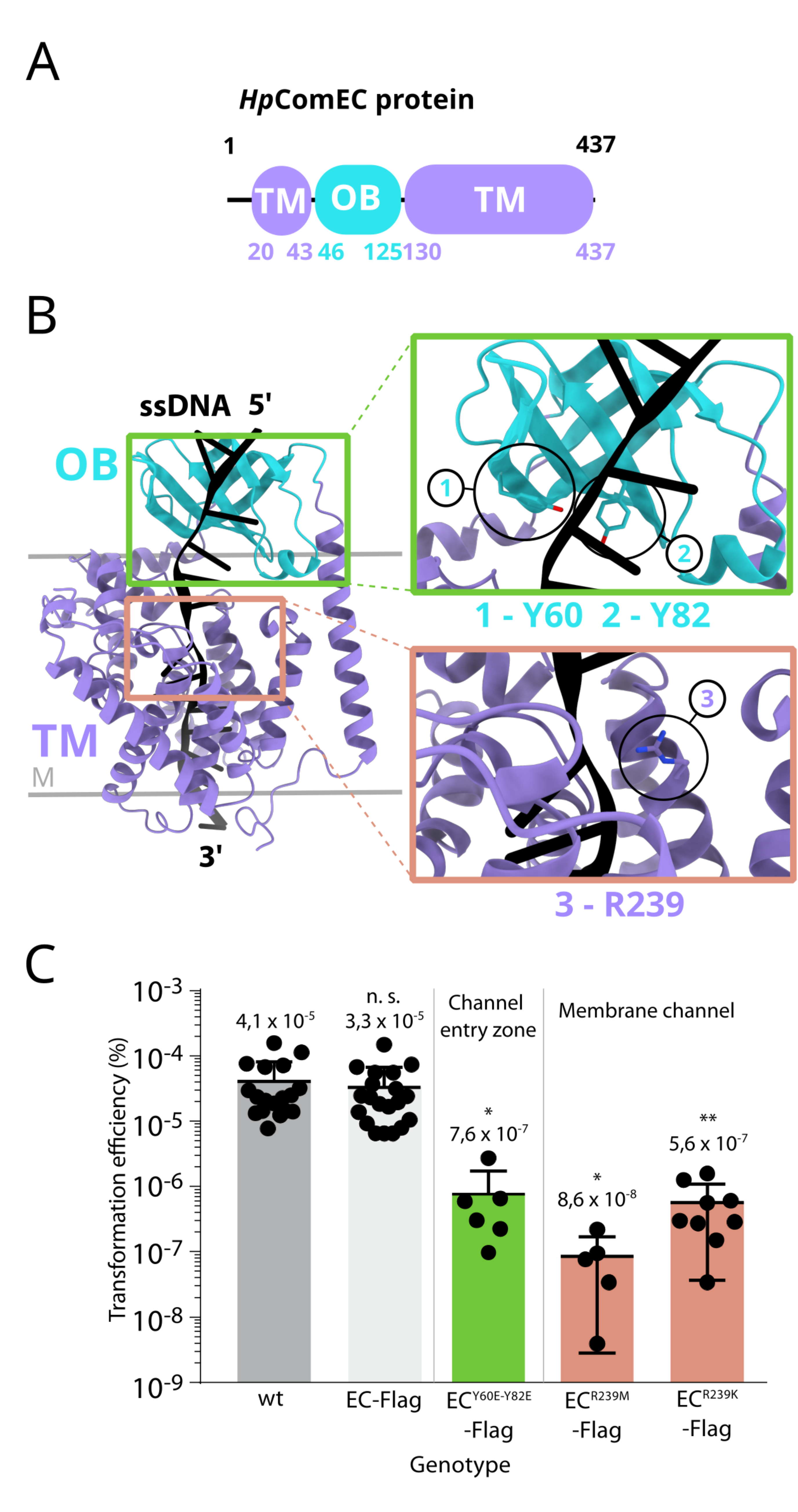
Mutation of selected residues on *Hp*ComEC protein from *Helicobacter pylori* affects transformation efficiency. (A) Domain organisation of *Hp*ComEC protein showing delimitations between two transmembrane domains (TM) and an OB-fold (OB). Numbers shown represent amino acids (AA) along the 437 AA protein. (B) Structural model of *Hp*ComEC protein interacting with ssDNA, showing transmembrane domains (purple) and OB-fold (light blue). Green insert shows Y60 (1) and Y82 (2) on OB-fold, predicted to interact with transforming ssDNA near the entry point of the membrane channel. Orange insert shows R239 (3) in the transmembrane channel, predicted to interact with transforming ssDNA. DNA polarity is shown. (C) Transformation efficiency of *HpcomEC* mutants when transformed with saturating concentrations of tDNA (strain 26695, Sm^R^). Strains used: *wt*, 1; *EC_Hp_-FLAG*, 1141; *EC_Hp_ ^Y60E-Y82E^-FLAG*, 1387; *EC_Hp_ ^R239M^-FLAG*, 1385; *EC_Hp_ ^R239K^ -FLAG*, 1389. Error bars representative of at least five-fold repeats, with individual data points shown. Student’s t-tests were used to compare transformation efficiencies of mutant strains with wildtype. **, p < 0.01; *, p < 0.05; n. s., not significant.

### A positively charged groove on *Sp*ComFC guides transforming ssDNA into the cytoplasm

The tripartite structural model of the pneumococcal tDNA transport machinery predicts that once ssDNA passes through the *Sp*ComEC transformation pore, it sequentially interacts with *Sp*ComFA and *Sp*ComFC. These proteins are specifically produced during competence. By tracking the cellular levels of *Sp*ComFA and *Sp*ComFC, we showed that their expression peaked 10-15 minutes after competence induction by CSP addition (Figure S6A). *Sp*ComFA was proposed to be an ATP-dependent ssDNA translocase with structural homology to SF2 DEAD family helicases, interacting with ssDNA via Walker A and B motifs^28^. However, the region of *Sp*ComFA predicted to interact with ssDNA did not allow selection of residues in which mutation would affect interaction without potentially affecting overall protein folding or stability. We thus focused on the interaction between internalised ssDNA and ComFC, previously identified for the homologous *H. pylori* ComF protein^27^. The model revealed a conserved, positively charged groove on the surface of *Sp*ComFC along which ssDNA is predicted to slide (Figure S1), with two residues, *Sp*ComFC^K83^ and *Sp*ComFC^R156^, central to this groove (Figure 4A). To assess the importance of these residues for transformation, both were mutated to negatively charged aspartate (D). These mutants were produced at wildtype levels during competence (Figure S6B). In saturating or limiting tDNA transformation assays, a *Sp*ComFC^K83D^ mutant showed a significant ∼7- or ∼120-fold decrease in transformation efficiency compared to wildtype respectively, while a *Sp*ComFC^R156D^ mutant had no significant effect (Figure 4B). However, a *Sp*ComFC^K83D-R156D^ double mutant displayed a cumulative effect, with a ∼50- or ∼2000-fold decrease, respectively (Figure 4B), significantly lower than the *Sp*ComFC^K83D^ mutant alone. These results strongly suggest that ssDNA interacts with *Sp*ComFC via the identified positively charged groove and that the charges of the K83 and R156 residues are central to this interface. This interaction is crucial for optimal ssDNA import and thus transformation efficiency.

**Figure 4:**
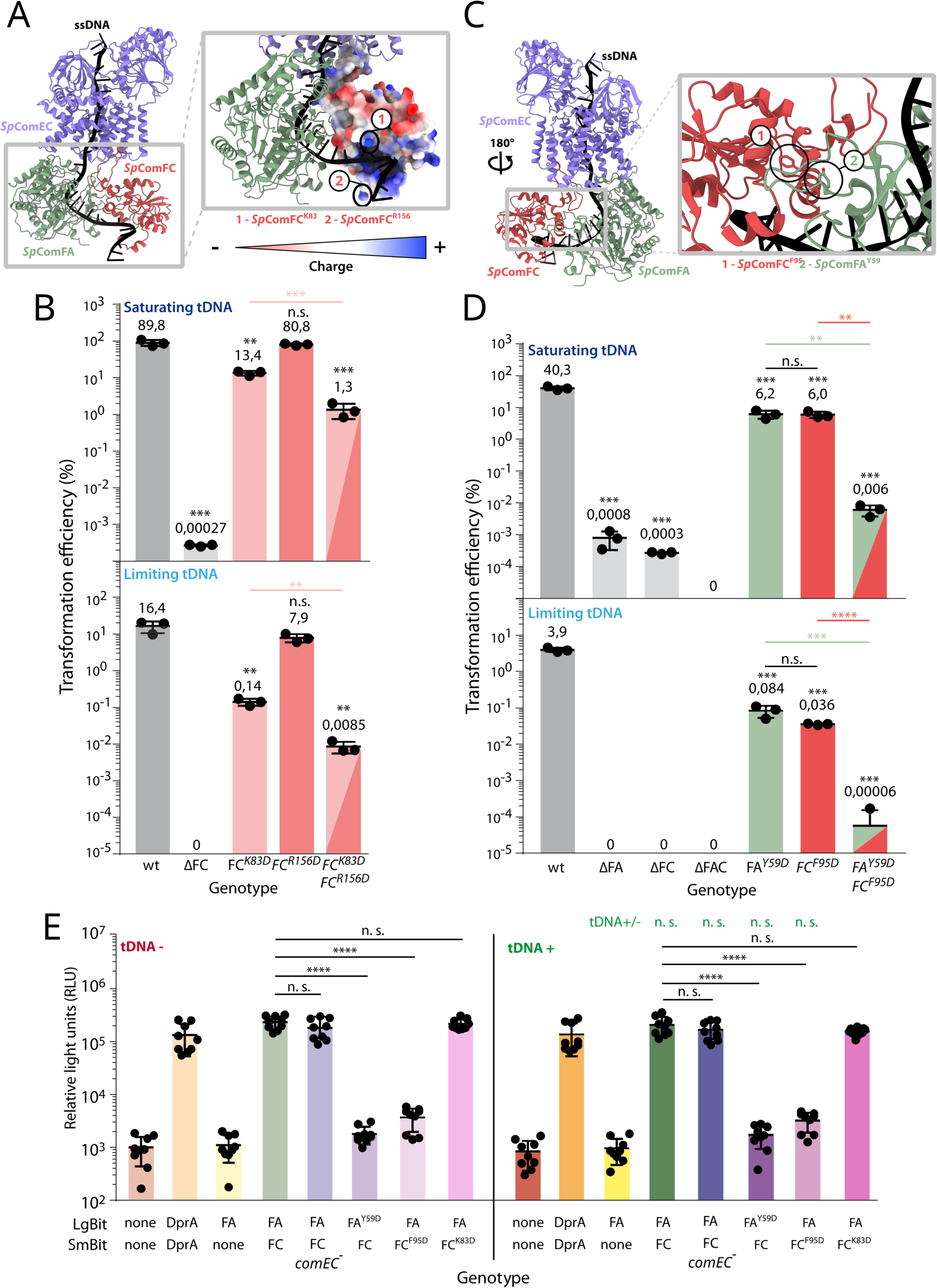
Interaction between *Sp*ComFC and tDNA or *Sp*ComFA is important for transformation. (A) Highlight of *Sp*ComFC-ssDNA interaction region on tripartite model of membrane transport complex. Electrostatic surface of *Sp*ComFC protein shown to highlight positively charged groove contacting ssDNA. Mutants predicted to affect interaction shown: (1) *Sp*ComFC^K83^; (2) *Sp*ComFC^R156^. (B) Transformation efficiency of single and double *Sp*ComFC mutants predicted to interact with ssDNA when transformed with saturating (2,500 pg mL^-1^) or limiting (25 pg mL^-1^) concentrations of tDNA. tDNA identity as in Figure 2F. Strains used: *wt*, R1501; *ΔFC*, R5099; *FC^K83D^*, R5191; *FC^R156D^*, R5238; *FC^K83D-R156D^*, R5270. Error bars representative of triplicate repeats, with individual data points shown. Student’s t-tests were used to compare transformation efficiencies of mutant strains with wildtype, unless shown by horizontal line. ***, p < 0.001; **, p < 0.01; n. s., not significant. (C) Highlight of *Sp*ComFA-*Sp*ComFC interaction region on tripartite model of membrane transport complex. Residues predicted as important for the interaction shown: (1) *Sp*ComFC^F95^; (2) *Sp*ComFA^Y59^. (D) Transformation efficiency of single and double pneumococcal ComFA/C mutants predicted to affect interaction between these proteins when transformed with saturating (2,500 pg mL^-1^) or limiting (25 pg mL^-1^) concentrations of tDNA. tDNA identity as in Figure 2F. Strains used: *wt*, R1501; *ΔFA*, R5114; *ΔFC*, R5099; *ΔFAC*, R4654; *FA^Y59D^*, R5064; *FC^F95D^*, R5153; *FA^Y59D^-FC^F95D^*, R5305. Error bars representative of triplicate repeats, with individual data points shown. Student’s t-tests were used to compare transformation efficiencies of mutant strains with wildtype, unless shown by horizontal line. ****, p < 0.0001; ***, p < 0.001; **, p < 0.01; n. s., not significant. (E) Split-luciferase assay of interaction between *Sp*ComFA and *Sp*ComFC. Strains used: red, R1501; orange, R4858; yellow, R5194; green, R5162; blue, R5252; indigo, R5179; violet, R5240; pink, R5247. Error bars representative of nine repeats, with means and individual data points shown. Student’s t-tests were used to compare luminescence of *Sp*ComFA/FC mutants with wildtype split luciferase strain as shown by horizontal lines. Green analyses compare tDNA+/-conditions. ****, p < 0,0001; n. s., not significant. Observed differences in the expression of ComFA and ComFC, along with transformation efficiency in these strains (Figure S6EF), is further discussed in the Supplementary Information section.

### Interaction between *Sp*ComFA and *Sp*ComFC is key for efficient pneumococcal transformation

Thus far, we identified several residues proposed to be key for interaction of ComEC and *Sp*ComFC with ssDNA. Next, we investigated the pairwise protein-protein interactions predicted in the tripartite model of ssDNA uptake in the pneumococcus, beginning with *Sp*ComFA and *Sp*ComFC. We first confirmed previous results showing that *Sp*ComFA and *Sp*ComFC are crucial for transformation^19,28^. Although residual transformants were observed when either protein remained present under saturating tDNA conditions, no transformants were observed when both proteins were absent (Figure 4D). An interaction between *Sp*ComFA and *Sp*ComFC has previously been documented in bacterial two-hybrid assays^28^. These two proteins are also proposed to interact directly in the tripartite model (Figure 1). Our model also predicted a strong interaction between these proteins (Table S2), and we identified two residues, *Sp*ComFA^Y59^ and *Sp*ComFC^F95^, potentially critical for this interaction (Figure 4C). Mutating each of these residues to aspartate (D) introduced a negative charge while preserving protein stability (Figure S6BC). Transformation assays under saturating tDNA conditions revealed that both *Sp*ComFA^Y59D^ and *Sp*ComFC^F95D^ mutations caused a significant 6-fold reduction in transformation efficiency compared to wildtype, showing that these mutations affect transformation, presumably by interfering with the interaction between *Sp*ComFA and *Sp*ComFC (Figure 4D). Accordingly, the double *Sp*ComFA^Y59D^-*Sp*ComFC^F95D^ mutant had a cumulative effect on transformation efficiency (Figure 4D), reducing transformants by ∼6,500-fold. To further assess the interaction between *Sp*ComFA and *Sp*ComFC in live pneumococcal cells, a split luciferase assay was carried out, where proximity of *Sp*ComFA-LgBit and SmBit-*Sp*ComFC in competent cells would reconstitute a luciferase protein and produce light^17,37,38^. Results show that a strain expressing *spcomFA-lgbit* alone displayed the same level of luminescence as a wildtype strain lacking either fusion (Figure 4E). However, a strain possessing both *spcomFA-lgbit* and *smbit-spcomFC* showed 200-fold higher luminescence, similar to that of a positive control strain possessing DprA dimers made up of DprA-LgBit and DprA-SmBit^16,17^ (Figure 4E). This result confirmed an interaction between *Sp*ComFA and *Sp*ComFC in live cells. This interaction was maintained in a Δ*spcomEC* background, showing that these two proteins interact even when the full tripartite complex is not formed. Both *Sp*ComFA^Y59D^ and *Sp*ComFC^F95D^ mutants display significantly lower luminescence levels (Figure 4E), confirming that these mutations disrupt the *Sp*ComFA-*Sp*ComFC interaction *in vivo*. As a control, we included the ComFC^K83D^ mutant, which affects interaction between ComFC and ssDNA (Figure 4B), but is not predicted to affect the ComFA-ComFC interaction. Accordingly, this mutant did not alter luminescence (Figure 4E). Addition of tDNA did not alter these results (Figure 4E), showing that the formation of this complex in competent cells is not dependent on active transformation. These results reveal residues key for the interaction between *Sp*ComFA and *Sp*ComFC, and show that this interaction is crucial for transformation, although it can occur independently of transformation.

### Exploring the importance of an interaction between *Sp*ComEC and *Sp*ComFA for pneumococcal transformation

A small interaction surface between *Sp*ComFA and a short cytoplasmic loop of *Sp*ComEC was predicted in the tripartite pneumococcal AlphaFold3 model (Figure 1 and Table S2). Three residues were predicted as potentially important for this interaction: *Sp*ComFA^W273^, *Sp*ComEC^R261^ and *Sp*ComEC^E267^ (Figure 5A). Substitution of the large, aromatic *Sp*ComFA^W273^ residue with a smaller residue, either neutral (alanine, A) or negatively charged (aspartate, D), did not affect transformation efficiency (Figure 5B). In addition, the *Sp*ComEC^R261A^ mutant and the double *Sp*ComFA^W273D^ *Sp*ComEC^R261A^ mutant did not affect transformation efficiency (Figure 5B). In contrast, a *Sp*ComEC^E267K^ mutant significantly reduced transformation efficiency (Figure 5B). All mutant proteins were stably produced (Figure S6CD). These results suggest that although there may be an interaction between *Sp*ComEC and *Sp*ComFA as part of the transforming ssDNA uptake complex, it is likely to be weak or transient.

**Figure 5:**
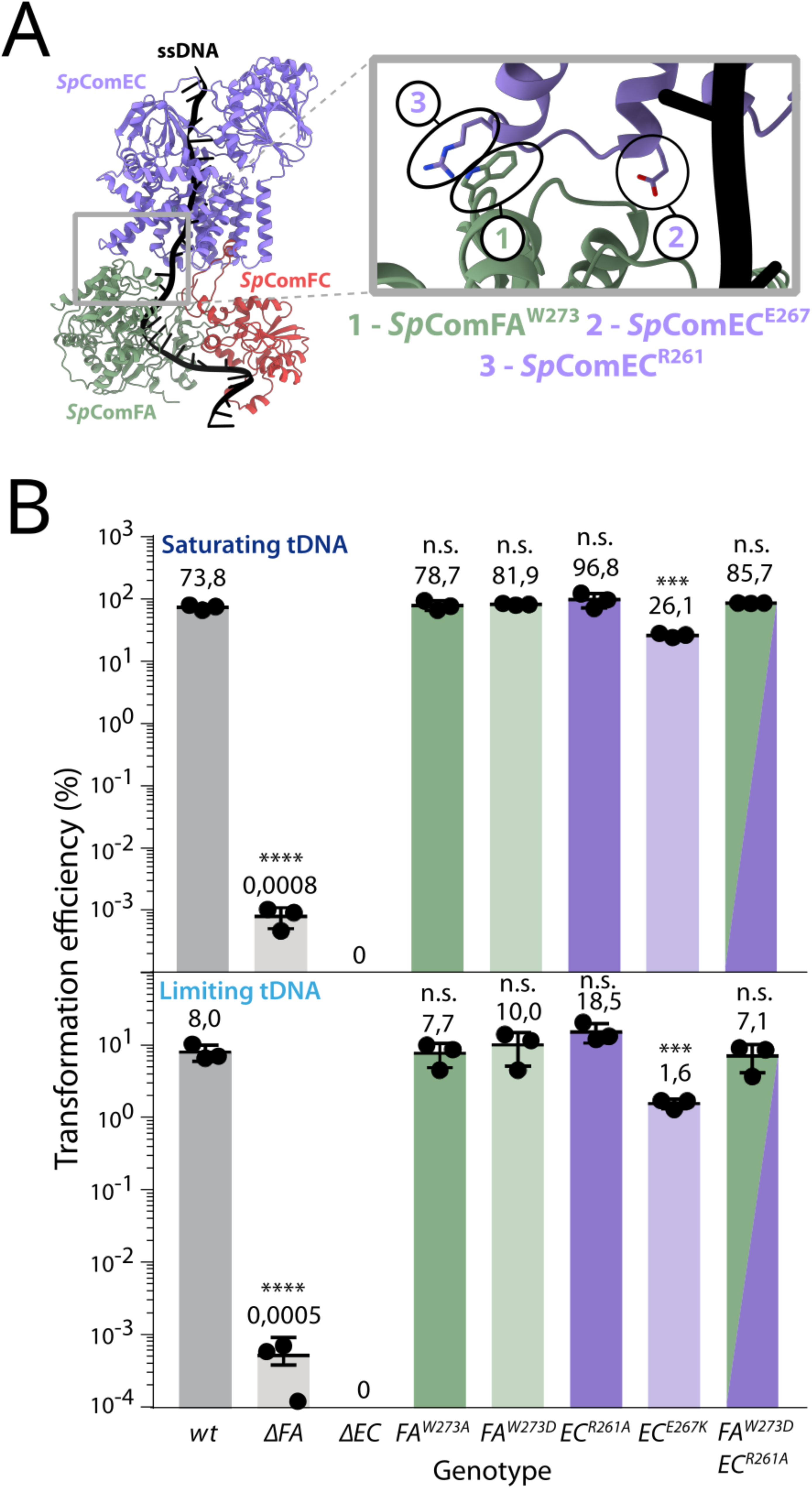
Interaction between *Sp*ComEC and *Sp*ComFA plays a minor role in transformation. (A) Highlight of *Sp*ComEC-*Sp*ComFA interaction region on tripartite model of membrane transport complex. Residues predicted as important for the interaction shown: (1) *Sp*ComFA^W273^; (2) *Sp*ComEC^E267^; (3) *Sp*ComEC^R261^. (B) Transformation efficiency of mutants predicted as important for interaction between *Sp*ComEC and *Sp*ComFA when transformed with saturating (2,500 pg mL^-1^) or limiting (25 pg mL^-1^) concentrations of tDNA. tDNA identity as in Figure 2F. Strains used: *wt*, R1501; *ΔFA*, R5114; *ΔEC*, R2300; *FA^W273A^*, R5278; *FA^W273D^*, R5367; *EC^R261A^*, R5281; *EC^E267K^*, R5282; *FA^W273D^, EC^R261A^*, R5368. Error bars representative of triplicate repeats, with individual data points shown. Student’s t-tests were used to compare transformation efficiencies of mutant strains with wildtype, unless shown by horizontal line. ****, p < 0.0001; ***, p < 0.001; n. s., not significant.

### A strong interaction between ComEC and ComFC is central for pneumococcal transformation and conserved in firmicutes

In the model of the tDNA uptake machinery in the pneumococcus, the interaction between *Sp*ComEC and *Sp*ComFC was characterized by the highest number of contacts and the largest contact area (Table S2). This interaction involves a small α-helix in the N-terminal part of *Sp*ComFC, which forms a ‘hook’ embedded within a ‘pocket’ in the *Sp*ComEC protein (Figure 6A). Two key residues were identified at this interface, namely aromatic *Sp*ComFC^F16^, on the *Sp*ComFC hook, and negatively charged *Sp*ComEC^D457^, in the *Sp*ComEC pocket (Figure 6A). These were replaced with aspartate (D) and lysine (K) residues, respectively. In addition, residues 12 to 26 of *Sp*ComFC were replaced with a flexible four-residue sequence (GSGG) to determine the effect of deleting the hook region on *Sp*ComEC-*Sp*ComFC interaction; this mutant is reported as *Sp*ComFC^Δ12-26^. All three mutants were stably produced in competent cells (Figure S6BD). In transformation assays with saturating tDNA, the *Sp*ComFC^Δ12-26^ strain displayed a dramatic >2,000-fold reduction in transformation efficiency compared to wildtype, showing the importance of the *Sp*ComFC hook for transformation (Figure 6B). Supporting this, the *Sp*ComFC^F16D^ mutation reduced transformation efficiency by ∼40-fold (Figure 6B). In addition, the *Sp*ComEC^D457K^ mutation displayed a significant 2-fold deficit in transformation efficiency (Figure 6B). Strikingly, the double *Sp*ComFC^F16D^ *Sp*ComEC^D457K^ mutant, which introduced complementary charge-swapping mutations at both sides of the interface, partially restored transformation efficiency compared to the *Sp*ComFC^F16D^ mutant (Figure 6B). This compensatory effect provides strong evidence that these two amino acids form a critical interaction pair within the *Sp*ComEC-*Sp*ComFC interface, consolidating the tripartite model. In transformation assays under limiting tDNA conditions, similar but more pronounced results were observed, strengthening the importance of this interaction for efficient tDNA uptake (Figure 6C).

**Figure 6:**
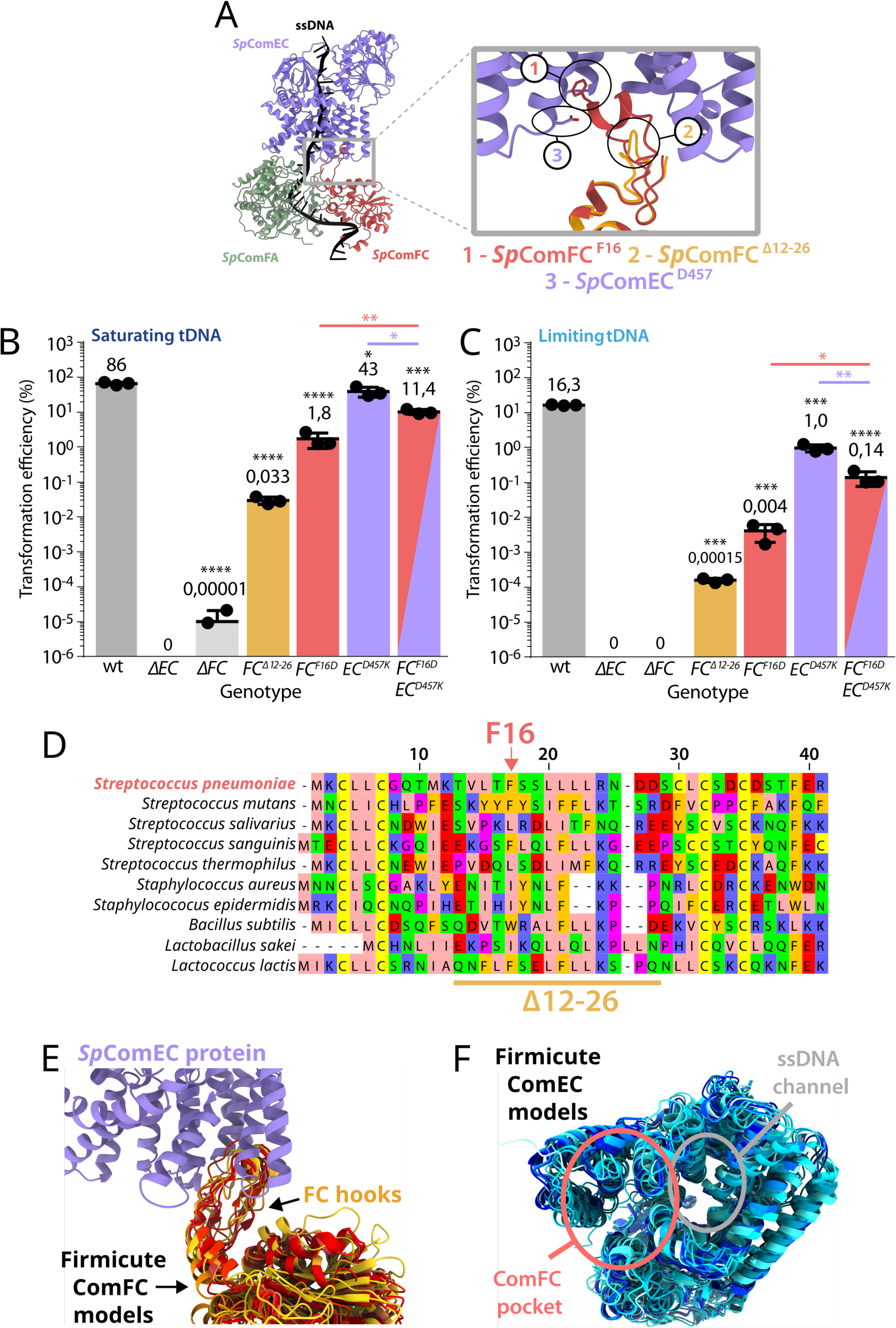
An interaction between *Sp*ComEC and *Sp*ComFC is key for pneumococcal transformation and structurally conserved in firmicutes. (A) Highlight of *Sp*ComEC-*Sp*ComFC interaction region on tripartite model of membrane transport complex. Highlight shows interaction zone of *Sp*ComFC ‘hook’ inserted into *Sp*ComEC ‘pocket’. Mutants predicted to affect interaction shown: (1) *Sp*ComFC^F16^, red; (2) *Sp*ComEC^D457^, purple. Structural model of deletion of *Sp*ComFC hook (Δ12-26) shown in orange (3). The highlighted zone is slightly pivoted compared to the full structure to facilitate observation of mutants. (B) Transformation efficiency of *Sp*ComEC/FC interaction mutants when transformed with saturating concentrations of tDNA. tDNA identity and concentration as in Figure 2F. *wt*, R1501; *ΔEC*, R2300; *ΔFC*, R5099; *FC^Δ12-26^*, R5280; *FC^F16D^*, R5279; *EC^D457K^*, R5283; *FC^F16D^ EC^D457K^*, R5366. Error bars representative of triplicate repeats, with individual data points shown. Student’s t-tests were used to compare transformation efficiencies of mutant strains with wildtype, unless shown by horizontal line. ****, p < 0.0001; ***, p < 0.001; *, p < 0.05. (C) Transformation efficiency of *Sp*ComEC/FC interaction mutants when transformed with limiting concentrations of tDNA. tDNA identity and concentration as in Figure 2G. Strains used, data representation and statistical analyses as in panel B. (D) AA alignment of *Sp*ComFC ‘hook’ region in selected firmicutes, with *S. pneumoniae* sequence and mutations (F16, Δ12-26) highlighted. Alignment Zappo AA colours as follows: blue, positive charge; red, negative charge; magenta, conformationally special; green, hydrophilic; pink, hydrophobic/aliphatic; orange, aromatic; yellow, cysteines. (E) Structural model alignment of ComFC ‘hook’ region in firmicute species from panel D, displayed with *Sp*ComEC (purple). Strains represented in order in a gradient from red to yellow: *S. pneumoniae, S. mutans, S. salivarius, S. sanguinis, S. thermophilus, S. aureus, S. epidermidis, B. subtilis, L. sakei, L. lactis.* (F) Structural alignment of ComEC in firmicute species from panel D, highlighting conservation of an ssDNA channel and a ‘pocket’ surrounded by α–helices into which the ComFC ‘hook’ is inserted. Strains represented in order in a gradient from dark to light blue: *S. pneumoniae, S. mutans, S. salivarius, S. sanguinis, S. thermophilus, S. aureus, S. epidermidis, B. subtilis, L. sakei, L. lactis*.

Since the interaction between *Sp*ComEC and *Sp*ComFC appeared key for transformation in the pneumococcus, we explored its structural conservation across a panel of bacterial species. The hook region was exclusively conserved in firmicutes (Table S1), mirroring the distribution of ComFA and suggesting that this ‘hook/pocket’ mode of interaction between ComEC and ComFC could be specific to species in which the tripartite complex ensures uptake of transforming ssDNA. Interestingly, although the hook region of *Sp*ComFC extends between two pairs of largely invariant cysteines, the amino acids in the hook itself are not strongly conserved (Figure 6D). However, aligning structural models of the ComEC-ComFC interaction showed that this ComFC hook extended into ComEC in all 10 firmicute species tested, suggesting that the interaction mode could be conserved despite amino acid variability (Figure 6E). In addition, alignment of ComEC models showed strong conservation of the ComEC pocket within which the ComFC hook inserts (Figure 6F). Taken together, these results suggest that this hook-pocket interaction between ComFC and ComEC is crucial for the integrity of the tripartite complex and thus for transformation in firmicutes.

### Exploring the effect of rupturing multiple interactions in the tripartite model on pneumococcal transformation

Transformation assays have confirmed the importance of specific interactions between pairs of proteins in the tripartite model of pneumococcal tDNA import. However, given the presence of multiple interaction surfaces between macromolecules in this model, we expect that the overall assembly could be robust with respect to the disruption of a single interface. Coupling of specific mutations important for each individual interaction should result in further destabilization of the tripartite tDNA import complex, and thus in greater impacts on transformation efficiency. To test this, a single mutation was chosen for each protein-protein interaction in the model: *Sp*ComFA^Y59D^ for *Sp*ComFA-*Sp*ComFC, *Sp*ComFA^W273A^ for *Sp*ComEC-*Sp*ComFA, *Sp*ComFC^F16D^ for *Sp*ComEC-*Sp*ComFC (Figure 7AB). Transformation efficiency of each single and double mutant was compared to wildtype in saturating and limiting tDNA transformation assays (Figure 7CD). As presented previously, *Sp*ComFA^Y59D^ and *Sp*ComFC^F16D^ single mutants displayed ∼6-fold and ∼40-fold decreases in transformation compared to wildtype, while *Sp*ComFA^W273A^ transformed at wildtype levels. In stark contrast, double mutants had drastic cumulative effects on transformation (Figure 7CD), despite mutant proteins being produced at wildtype levels during competence (Figure S7ABC). In addition, the *Sp*ComFC^F16D^ mutant was also combined with another mutation predicted to affect interaction between *Sp*ComEC and *Sp*ComFA, but showing no decrease in transformation efficiency (*Sp*ComEC^R261A^). Again, this combination led to a significant cumulative decrease in transformation efficiency (Figure 7CD). These striking results showed that affecting two interactions within the tripartite complex has drastic effects on transformation, even when one of these mutations alone (*Sp*ComFA^W273A^ or *Sp*ComEC^R261A^) had no effect on transformation efficiency. This suggests that the interaction between *Sp*ComEC and *Sp*ComFA can play a stabilising role in the complex when the strong interactions of *Sp*ComFC with either *Sp*ComFA (*Sp*ComFA^Y59D^) or *Sp*ComEC (*Sp*ComFC^F16D^) are affected. These cumulative effects provide strong experimental validation for the tripartite model of tDNA import.

**Figure 7:**
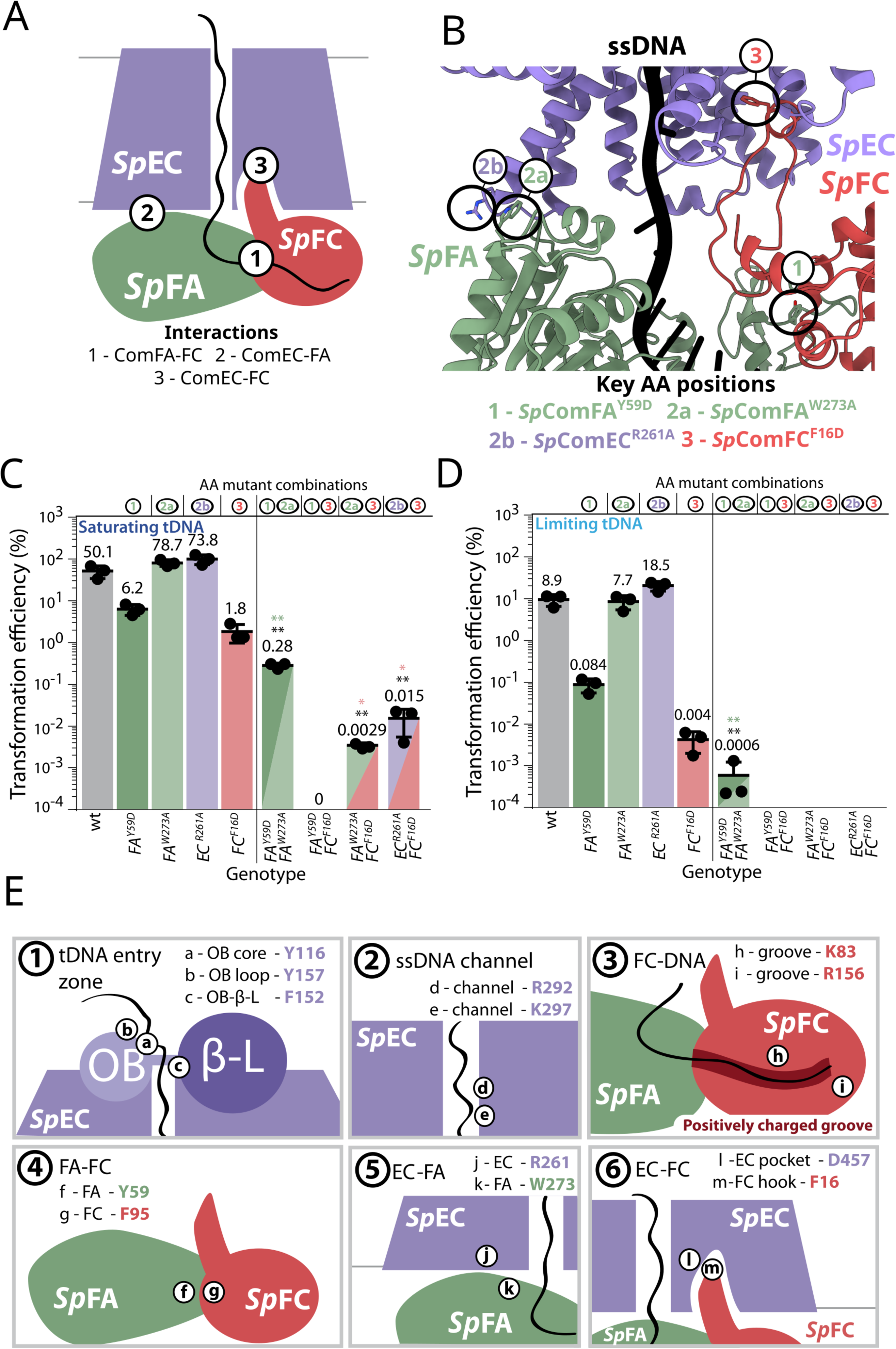
Double mutations targeting different interactions in the tripartite *Sp*ComEC/FC/FA model have cumulative effects on transformation. (A) Schematic representation of tripartite model of tDNA membrane transport, showing three key protein-protein interaction zones. (B) Highlight of tripartite structural model of tDNA membrane transport, with three residues shown, one key to each protein-protein interaction tested for this model: (1) *Sp*ComFA^Y59D^; (2) *Sp*ComFA^W273A^; (3) *Sp*ComFC^F16D^. (C) Transformation efficiency of single or double interaction mutants when transformed with saturating concentrations of tDNA. tDNA identity and concentration as in Figure 2F. Strains used: *wt*, R1501; *FA^Y59D^*, R5064, *FA^W273A^*, R5278; *EC^R261A^*, R2581; *FC^F16D^*, R5279; *FA^Y59D-W273A^*, R5395; *FA^Y59D^ FC^F16D^*, R5309; *FA^W273A^ FC^F16D^*, R5307; *FC^F16D^ EC^R261A^*, R5393. Error bars representative of triplicate repeats, with individual data points shown. (D) Transformation efficiency of double interaction mutants when transformed with limiting concentrations of tDNA. tDNA identity and concentration as in Figure 2G. Data representation as in panel C. (E) Schematic zoom of six interaction zones on the tripartite complex of tDNA membrane transport during transformation in firmicutes, highlighting important residues identified in each interaction in this study.

To assess whether protein-tDNA interactions are similarly important, we combined the *Sp*ComFA^Y59D^ and *Sp*ComFC^F16D^ mutations with the *Sp*ComFC^K83D^ mutation, predicted to disrupt the interaction of a positively charged channel on *Sp*ComFC with tDNA (Figure S8AB). The combination resulted in a ∼1,000-fold decrease in transformation efficiency under saturating tDNA conditions and only residual transformants observed in limiting tDNA conditions (Figure S8CD), despite mutant proteins being produced at wildtype levels during competence in these strains (Figure S7A).

Taken together, these results demonstrate that combining mutations affecting individual interactions within the tripartite structural model of pneumococcal tDNA uptake has strong cumulative effects. This provides strong support for this model, revealing for the first time that the interactions between *Sp*ComEC, *Sp*ComFA and *Sp*ComFC are crucial for the import of transforming ssDNA. We propose that this tripartite complex constitutes a central and conserved mechanism for transformation across firmicutes.

## Discussion

Historically, the study of transmembrane proteins has been hindered by the difficulty in obtaining accurate structural information^39,40^. New developments in AI-based structural modelling software such as AlphaFold^41^ have enabled the accurate structural modelling of transmembrane proteins^42^ and protein-protein complexes^43^. The most recent version, AlphaFold3^31^, enables modelling of complexes including proteins and nucleic acids, albeit with an overall lower success rate for protein-nucleic acid interactions compared to protein-protein interactions. Here, we leveraged these recent advances to study the mechanism of ssDNA import across the bacterial membrane during natural transformation. Specifically, we focused on the interaction of the ComEC transmembrane channel protein with the cytoplasmic ComFA/ComFC complex^28^ and with ssDNA. The molecular mechanisms governing this process have remained elusive due to the difficulty in working with ComEC. Using structural models supported by *in vivo* data, we show that in the human pathogen *S. pneumoniae*, *Sp*ComEC forms a tripartite complex with *Sp*ComFA and *Sp*ComFC to perform import of ssDNA across the membrane during transformation, and that this complex is structurally conserved in firmicutes. AlphaFold3 modelling played several important roles in this study, firstly as a hypothesis generator identifying possible interactions and key residues, secondly as a way to assess evolutionary conservation of complex structures, and finally as a tool for designing experiments, e.g. in identifying a cytoplasmic loop of *Sp*ComEC for alfa-tag insertion with minimal impact on structure, which enabled the monitoring of mutant protein expression levels (Figure S3A). Overall, this study thus provides a strong proof of principle of the use for AlphaFold3 to probe the role of macromolecular interactions in biological processes, based on a strong readout of altered interactions, in this case the selection of an antibiotic resistance marker integrated by transformation.

*Sp*ComEC possesses three domains, a transmembrane *Competence* domain as well as an OB-fold and a β-lactamase domain, both situated in the periplasmic space in our model (Figure 2). Although the *Competence* domain is conserved across transformable species, the other two domains are not^20^. Here we show that the pneumococcal OB-fold interacts both with the β-lactamase domain and with transforming ssDNA (Figure 7E, box 1), in line with previous *in silico* studies proposing the *B. subtilis* OB-fold as binding ssDNA^21^. A recent preprint investigated the structural organization and activity of the DNA-binding OB-fold domain and β-lactamase (nuclease) domain of ComEC in another firmicute species (*Moorella glycerini*)^23^. The preprint proposes that the two domains are key for simultaneous achievement of degradation of one DNA strand and translocation of the other strand through the ComEC channel. Some of the mutations tested in that study in *Legionella pneumophila* confirm our results on the importance of channel entry residues for ssDNA translocation. More generally, how ComEC interfaces with key components of the transformation machinery on the outside of the membrane, such as ComEA and EndA, remains to be established. The outward-facing OB-fold and β-lactamase domain may be involved in such interactions. In line with this idea, *Hp*ComEC interacts through its OB-fold domain with ComH, a periplasmic protein functionally analogous to ComEA^44^.

In all firmicute models, the ssDNA passes through the conserved ComEC channel with the 3’ end entering the cell first. This is in agreement with historical genetic studies in the pneumococcus^26,34^ and a recent study in other firmicute species^23^, and in contrast with a recent suggestion of opposite polarity based on 5’-3’ polarity of *Sp*ComFA translocation on ssDNA in *in vitro* assays^29^. We validated the importance of two residues within the *Sp*ComEC channel (Figure 7E, box 2). The conservation of the transmembrane channel structure was demonstrated by the importance of a conserved arginine residue in transformation for both monoderm (R292 in *Sp*ComEC, Figure 2) and diderm (R239 in *Hp*ComEC, Figure 3) species. This strongly suggests that although the accessory domains (OB-fold, β-lactamase) may differ between species, the membrane channel-forming *Competence* domain is structurally conserved across transformable bacterial species.

The tripartite protein models generated by AlphaFold3 revealed a conserved interaction mode of *Sp*ComEC, *Sp*ComFA and *Sp*ComFC in firmicutes. The pneumococcal model allowed identification of residues key for several interactions in the model, which were validated by assaying the effect of mutating these residues on transformation. Through this method, we revealed a groove on *Sp*ComFC along which ssDNA slides, and residues important for interactions between *Sp*ComFA and *Sp*ComFC, *Sp*ComEC and *Sp*ComFA and *Sp*ComEC and *Sp*ComFC (Figure 7E, boxes 3-6 respectively). Although several residues predicted to affect *Sp*ComEC-*Sp*ComFA interaction had no effect on transformation when mutated alone, coupling mutation of residues predicted to be important for the *Sp*ComEC-*Sp*ComFA interaction with those key for *Sp*ComEC-*Sp*ComFC or *Sp*ComFA-*Sp*ComFC interactions had a cumulative effect on transformation efficiency. This strongly suggests that the predicted *Sp*ComEC-*Sp*ComFA interaction plays a role in stabilising the tripartite complex, but that this is only revealed when simultaneously disrupting another pairwise interaction in the tripartite complex. Indeed, the fact that all of the mutants tested were found to have cumulative effects on transformation (Figures 7 and S8) provides strong validation of this model.

A ‘hook and pocket’ interaction crucial for transformation was revealed between *Sp*ComFC and *Sp*ComEC, where an α-helix in the N-terminal of *Sp*ComFC, flanked by conserved cysteine bridges, protrudes into the *Sp*ComEC protein (Figure 6). Despite the presence of ComFC in all transformable species, the hook structure was conserved only in firmicutes (Table S1). However, previously shown specific binding of *Hp*ComF with ssDNA and dependence of *Hp*ComF association with the membrane on the presence of *Hp*ComEC^27^ indicate that the ComEC-ComF(C)-ssDNA interaction may be generalised, albeit through distinct mechanisms. More generally, the central role of ComEC as an interaction platform essential to natural transformation seems to be universal, despite possibly different interaction partners and interaction modes, on both sides of the membrane.

Using AlphaFold3 modelling, we were unable to identify residues central to the interaction between *Sp*ComFA and ssDNA but not affecting *Sp*ComFA stability. However, *Sp*ComFA has been shown to have ssDNA-dependent ATPase activity *in vitro*^28^, with this activity proposed to pull ssDNA through the *Sp*ComEC channel. Accordingly, mutations in the conserved Walker A or B boxes of the ATPase domain impacted transformation in *B. subtilis*^45^, and a Walker B mutant reduced ATPase activity of pneumococcal ComFA *in vitro*^28^. As with the ComFC ‘hook’ motif, ComFA is conserved only in firmicutes (Table S1). The structural conservation of the tripartite protein model (Figure S2) suggests that the role of *Sp*ComFA as a motor for ssDNA import may be conserved across firmicutes. Once internalised, ssDNA interacts with conserved proteins involved in homologous recombination, including the homologous recombinase RecA and the transformation-dedicated loader DprA^15^. Although the interactions between proteins at the frontier of ssDNA import and homologous recombination remain poorly defined, ComFA, RecA and DprA colocalise at the cell pole of *B. subtilis* cells^46,47^, while an interaction between *Sp*ComFA and DprA was observed in bacterial two hybrid assays^28^. Further study is required to reveal the nature of the interactions ensuring this next step in the transformation process.

In conclusion, our study has coupled structural modelling using AlphaFold3 with functional assays to reveal a tripartite protein complex made up of the transmembrane channel *Sp*ComEC, the ATPase *Sp*ComFA and the ssDNA-binding protein *Sp*ComFC. This approach allowed circumvention of the issues of working with the transmembrane *Sp*ComEC protein, and revealed key interactions and residues important for ssDNA import across the membrane during transformation. In addition, alignment of protein sequences and structural models highlighted that this tripartite model was conserved in firmicutes. As well as validating this approach for the study of macromolecular complexes, including those interacting with DNA, this study reveals and validates for the first time how three proteins coordinate together to ensure the import of ssDNA into cells during transformation. This provides a previously unattainable understanding of the molecular mechanisms involved in ssDNA import in this key horizontal gene transfer mechanism, which is central to bacterial adaptation.

## Materials and Methods

### *S. pneumoniae* strains, competence and transformation

The pneumococcal strains and primers used in this study can be found in (Table S3). Standard procedures for transformation and growth media were used^48^. In this study, cells were prevented from spontaneously developing competence by deletion of the *comC* gene (*comC0*)^5^. Pre-competent cultures were prepared by growing cells to an OD_550_ of 0.1 in C+Y medium (pH 7) before 10-fold concentration and storage at –80°C as 100 μL aliquots and transformation was carried out as previously described with modifications^49^. Briefly, in assays to compare transformation efficiency, 100 μL aliquots of pre-competent cells were resuspended in 900 μL fresh C+Y medium with 100 ng mL^−1^ CSP and incubated at 37°C for 10 min. 5 ng µL^-1^ transforming DNA (saturating condition) or 0.05 ng µL^-1^ transforming DNA (limiting condition) was then added to a 100 μL aliquot of this culture, followed by incubation at 30°C for 20 min. Cells were then diluted and plated on 10 mL CAT agar with 5% horse blood before incubation at 37°C for 2 hr. A further 10 mL layer of CAT agar with appropriate antibiotic was added to plates to select transformants, and plates without antibiotic were used as comparison to calculate transformation efficiency where appropriate. Plates were incubated overnight at 37°C. To compare transformation efficiencies, transforming DNA was a 3,434 bp PCR fragment amplified with primer pair MB117-MB120 as noted^35^, possessing a *rpsL41* point mutation conferring streptomycin resistance. Antibiotic concentrations (μg mL^−1^) used for the selection of *S. pneumoniae* transformants were spectinomycin (Spc), 100; streptomycin (Sm), 200; and trimethoprim (Trim), 20. GraphPad Prism was used for statistical analyses. To compare two averages of triplicate repeats, an unpaired *t* test was used. Detailed information regarding the construction of new strains can be found in the Supplementary Information section.

### *H. pylori* growth conditions

All *H. pylori* strains used in this study are derivatives of strain 26695^50^. Cultures were grown at 37°C under microaerobic conditions (5 % O_2_, 10 % CO_2_), using the Whitley Jar Gassing System (Don Whitley Scientific), and 100 % humidity. For plate cultures, a blood agar base (BAB) or Brucella medium enriched with 10 % defibrillated horse blood (ThermoFisher SR0050) was employed, supplemented with an antibiotic mix containing polymyxin B (0.155 mg mL^-1^), vancomycin (6.25 mg mL^-1^), trimethoprim (3.125 mg mL^-1^), and amphotericin B (1.25 mg mL^-1^). Additional antibiotics were added as required: kanamycin (20 µg mL^-1^), chloramphenicol (8 µg mL^-1^), and streptomycin (10 μg mL^-1^).

### Natural transformation of *H. pylori*

Introduction of the desired constructs by natural transformation into *H. pylori* strains was performed as previously described^51^. Briefly, 15 µL (optical density of 4.0 at 600 nm) of exponentially *H. pylori* culture were spotted with plasmid DNA (200 ng) containing the desired construct on Brucella plates and incubated for 24 hours at 37 °C. Next day, spots were recovered, cells resuspended and plated on Brucella plates with either kanamycin (20 μg mL^-1^) or chloramphenicol (8 μg mL^-1^) and incubated for 4–5 days at 37 °C. The clones were subsequently checked by PCR or sequencing to confirm the presence of the deletion or verify the sequence of the expressed gene, respectively.

### Determination of natural transformation frequencies in *H. pylori*

Natural transformation assays were performed as described^44^. Briefly, 3 x 15 µL of exponentially growing *H. pylori* cells (optical density of 4.0 at 600 nm) were spotted incubated in the presence chromosomal DNA (200 ng) of streptomycin resistant 26695 *H. pylori* strain on Brucella plates for 24 h at 37 °C. Next day, the spots were recovered and the appropriate dilutions of *H. pylori* cells were plated on Brucella plates with or without streptomycin (10 μg mL^-1^) and incubated for 4–5 days at 37 °C. The frequency of natural transformation was determined by comparing the number of colonies obtained on media containing streptomycin to the number of colonies on media without streptomycin. P values were calculated using the Mann–Whitney U test on GraphPad Prism software.

### Western blot analysis

To determine the kinetics of *Sp*ComFA, *Sp*ComFC and *Sp*ComEC protein production during transformation, pneumococcal cells were grown in C+Y medium at 37°C to OD_550_ 0.1. 2.5 mL of culture was sampled (T0) and added to 2 mL of ice-cold C+Y to stop growth. 100 ng mL-1 CSP was added to the cells and 2.5 mL of culture was sampled as with the T=0 sample every 5 min for 20 min and then every 10 min for 70 min. Samples were centrifuged (10 min, 5000 g) and pellets were resuspended in 50 µL of TE 1x supplemented with 0.01% DOC and 0.02% SDS. Samples were then incubated for 10 min at 37°C before addition of 20 µL 5x sample buffer with 10% β-mercaptoethanol, followed by incubation at 95°C for 5 min. To compare protein expression profiles, Western blots were carried out as previously described^49^. Primary polyclonal antibodies raised against *Sp*ComFA and SsbB were used at 1/10,000 dilution. Primary monoclonal recombinant IgG antibodies raised against ALFA-tag (Nanotag Biotechnologies) were used at 1/3,000 dilution. Primary polyclonal antibodies raised against ComFC were used at 1/1,000 dilution.

For *H. pylori* samples, membrane protein extracts were prepared by sonicating cells with a Bioruptor bath (pulses 15s on and 15s off for 10 min at maximum intensity) in PBS buffer (PBS 1 X, 150 mM NaCl, 10% glycerol, 150 mM NaCl), with COMPLETE^TM^ ( Protease Inhibitor Cocktail Tablets from Merck Ref: 11836170001) and centrifugation for 20 min at 14,000 rpm at 4 °C. The supernatant was removed and set aside. The pellet was washed with PBS buffer and then suspended in PBS buffer by sonication with the Bioruptor bath (pulses 15 s on and 15 s off for 5 min at full intensity). The protein concentration of the membrane fraction was measured with the Bradford reagent (Sigma B6916) using a bovine serum albumin as standard. 100 µg of protein from the membrane fraction were heated in Laemmli for 15 min at 50 °C, then separated on a 4-15 % gradient SDS-PAGE gel, and transferred to a nitrocellulose membrane. The membrane was blocked with 5% milk in PBST (PBS 1 X; 0.3% Tween 20; 150 mM NaCl), and incubated with anti-FLAG (Sigma F1804-200UG) or anti-MotB (kind gift from Hilde De Reuse, Pasteur Institute) antibodies. Goat anti-rabbit 800 antibody (Diagomics R-05060-250) or goat anti-mouse 800 antibody (Diagomics R-05061-250) were used as secondary antibodies and blots were detected using the Cytivia Typhoon Biomolecular Imager.

### Split-Luciferase Assay

Split luciferase assays were carried out as previously described^37,38,52^, with modifications. Briefly, pneumococcal cells were grown in a C + Y medium (with 50 µM IPTG where required) at 37 °C until OD_550_ 0.1, and competence was induced by the addition of 100 ng mL^−1^ CSP. The cells were then incubated for 10 min at 37 °C before the addition of R1501 chromosomal DNA (250 ng µL^−1^) where noted, followed by a further 5 min incubation at 37 °C. The cells were then washed in fresh C + Y medium and 1% NanoGlo substrate (Promega) was added and luminescence was measured 20 times every 30 seconds in a plate reader (VarioSkan luminometer, ThermoFisher). Data are represented as mean maximal luminescence ± SD calculated from nine independent repeats, with individual data points plotted, as shown previously^17^.

### Structural analyses

All structural models were obtained using the AlphaFold3^31^ web server and visualized using the ChimeraX^53^ software. Cartoon representations were selected to show proteins and DNA except for residues of interest highlighted by a stick representation and coloured by heteroatoms. Electrostatic potential of the *Sp*ComFC protein surface was computed using the “electrostatic” ChimeraX module. Structural alignments were computed thanks to the “matchmaker” feature of ChimeraX using global alignment strategies with *S. pneumoniae* proteins as reference, keeping default parameters. RMSD values between firmicute AlphaFold3 models and the *S. pneumoniae* model were obtained using a pruning iteration cut-off distance of 5 Å (to exclude outlier atoms) and excluding DNA from these calculations.

Sequence identities between *S. pneumoniae* and other firmicutes were computed for each protein of the complex based on pairwise structural alignments, using RCSB PDB Pairwise Structural Alignment web server^54^. The input structures were *Sp*ComEC, *Sp*ComFC and *Sp*ComFA protein models derived from the AF3 quadruplex model.

Structural interaction analyses were made using ChimeraX “measure buriedarea” command to compute the exact interacting surface between ComEC/ComFC, ComEC/ComFA and ComFC/ComFA, using AF3 models as reference interaction structure. The “contact” feature was used on the same models to compute the number of contacts implied in the interaction using a distance threshold of 5.00 Å and other options as default.

### ComEC domain identification

Domains were first identified by querying The Encyclopedia of Domains (TED)^55^ using *Sp*ComEC and *Hp*ComEC UniProt^56^ IDs (respectively, Q8DQ40 and O25915). A total of 5 domains were predicted for *Sp*ComEC including the OB-fold and β-lactamase domain with associated CATH topology-level identifiers (respectively, 2.40.50 and 3.60.15). We merged the other 3 domains to form a single discontinuous transmembrane domain. For *Hp*ComEC, 3 domains were initially detected, including the OB-fold (CATH topology-level identifier 2.40.50). The other 2 domains and unassigned regions were fused to form the discontinuous transmembrane domain of the protein.

### ComFC sequence alignment in firmicutes

ComFC sequences were retrieved from their respective UniProt IDs (Table S2) identified by a keyword research over the UniProt database. They were then aligned using the MAFFT accuracy-oriented E-INS-i method^57^. Sequence alignment is visualized using Jalview^58^ and colorued based on the Zappo color scheme.

### Evolutionary conservation of *Sp*ComEC, *Hp*ComEC, *Sp*ComFA and *Sp*ComFC

For each protein, homologous sequences were retrieved using MMseqs2^59^ against the Uniref30^60^ version dated February 2022. Redundancy was filtered with HHfilter^61^, using 70% minimum sequence identity and a diff parameter of 100, limiting the number of sequences while ensuring sequence diversity in the MSA. Evolutionary conservation was mapped from the MSA onto the structural models using conservation scores computed with Rate4Site^62,63^. Visualization was done using ChimeraX b-factor coloration and a white to red color scale.

## Supporting information

Dewailly Fauconnet 2025 - Supplementary Tables

Dewailly Fauconnet 2025 - Supplementary Figures

Dewailly Fauconnet 2025 - Supplementary Information

## Acknowledgements

This work was supported by grant ANR-22-CE44-0044 from Agence Nationale de la Recherche to PP, JPR, JA and RG. We thank the BIOI2 platform for making the ColabFold and MSA-tools pipelines easily accessible at the I2BC for YF, JA and RG. This work was granted access to the HPC resources of IDRIS under the allocation 2024-AD010314343R1 by GENCI to RG.

