## Supplementary material for "A tripartite protein complex promotes DNA transport during natural transformation in firmicutes": Dewailly Fauconnet 2025 - Supplementary Tables

| Phylum | Species name | Gram | ComEC | ComF/ComFC | ComFA | ComFC N-term hook |
|---|---|---|---|---|---|---|
| **Gammaproteobacteria** | *Vibrio cholerae* | - | Q9KQW8 | Q9KNL3 | - | - |
| | *Haemophilus influenzae* | - | P44408 | P31773 | - | - |
| | *Acintetobacter baunamii* | - | A0A6F8TDY5 | A0A6F8TCP7 | - | - |
| | *Acintobacter baylyi* | - | Q9EYM1 | Q6F7P6 | - | - |
| | *Legionella pneumophila* | - | Q5ZXV5 | Q5ZT33 | - | - |
| **Betaproteobacteria** | *Neisseria gonorrhoeae* | - | Q5F9W1 | Q5F640 | - | - |
| **Firmicutes** | *Bacillus subtilis* | + | P39695 | P39147 | P39145 | + |
| | *Staphylococcus aureus* | + | A0A6K1MRM3 | A0A6B5I2H0 | A0A3A5M2J4 | + |
| | *Streptococcus sanguinis* | + | A3CLU7 | A3CPW2 | A3CPW3 | + |
| | *Streptococcus mutans* | + | Q8DV79 | Q8DVI9 | Q8DVJ0 | + |
| | *Streptococcus thermophilus* | + | A0A2S4DYQ8 | A0A2X3US27 | A0A2X3UVJ1 | + |
| | *Streptococcus salivarius* | + | A0A6N3BLK3 | A0A7L6WIW1 | A0A448A7C8 | + |
| | *Streptococcus pneumoniae* | + | Q8DQ40 | Q8CWN0 | Q8CWM9 | + |
| | *Lactobacillus sakei* | + | Q38WR1 | Q38YD4 | Q38YD5 | + |
| | *Lactococcus lactis* | + | Q9CER4 | Q9CGK7 | Q9CGK6 | + |
| | *Staphylococcus epidermidis* | + | Q5HNV9 | Q5HQX8 | Q5HQX9 | + |
| **Cyanobacteria** | *Thermosynechococcus elongatus* | - | A0A7D6EVL4 | A0A7D6J197 | - | - |
| **Bacteroidetes** | *Porphyromonas gingivalis* | - | A0A0E2LR21 | A0A0E2LU95 | - | - |
| **Chlorobi** | *Chlorobium tepidum* | - | Q8KCP6 | Q8KAZ2 | - | - |
| **Deinococcus** | *Thermus thermophilus* | - | Q5SGW2 | Q5SJE3 | - | - |
| **Enterobacteria** | *Salmonella enterica* | - | A0A3F3I8H2 | A0A1Z3X6N5 | - | - |
| | *Escherichia coli* | - | P37443 | P46846 | - | - |
| **Alphaproteobacteria** | *Agrobacterium tumefaciens* | - | A0A4D7YWA1 | A0A4D7Z1C4 | - | - |
| **Epsilonproteobacteria** | *Helicobacter pylori* | - | O25915 | O26008 | - | - |
| **Actinobacteria** | *Mycobacterium bovis* | + | A0AAE8WLB3 | A0AAE9BB63 | - | - |
| | *Streptomyces kasugaensis* | + | A0A4V2JHZ0 | A0A4Q9HPW3 | - | - |
| | *Corynebacterium glutamicum ATCC* | + | Q8NN62 | O87330 | - | - |
| | *Corynebacterium glutamicum R* | + | A4QG68 | O87330 | - | - |
| **Mycoplasma** | *Mycoplasma mycoides* | * | A0A654IKZ2 | Q6MTD4 | - | - |
| **Chlamydia** | *Chlamidia pneumoniae* | - | Q9JS51 | A0A0F7WSN5 | - | - |
| | *Chlamydia muridarum* | - | Q9PK55 | A0A070A0H0 | - | - |
| **Leptospira** | *Leptospira interogans* | - | Q8F8Z2 | Q8F8Q1 | - | - |
| **Spirochete** | *Treponema denticola SP33* | - | M2BJL0 | M2C3D6 | - | - |

* Bacterium without cell wall.

| | Uniprot ID | | | AF3 modelisation | | | Interacting surface area (Å²)** | | | Seq. identity*** (%) | | |
|---|---|---|---|---|---|---|---|---|---|---|---|---|
| **Species** | **ComEC** | **ComFC** | **ComFA** | **ipTM** | **pTM** | **RMSD*** | **EC/FC** | **EC/FA** | **FC/FA** | **EC** | **FC** | **FA** |
| ***Streptococcus pneumoniae*** | **Q8DQ40** | **Q8CWN0** | **Q8CWM9** | **0.74** | **0.82** | **0** | **816** | **366** | **535** | **-** | **-** | **-** |
| *Streptococcus sanguinis* | A3CLU7 | A3CPW2 | A3CPW3 | 0.76 | 0.83 | 2.14 | 865 | 443 | 590 | 56 | 46 | 70 |
| *Streptococcus mutans* | Q8DV79 | Q8DVI9 | Q8DVJ0 | 0.76 | 0.84 | 1.94 | 980 | 426 | 485 | 50 | 44 | 59 |
| *Streptococcus thermophilus* | A0A2S4DYQ8 | A0A2X3US27 | A0A2X3UVJ1 | 0.74 | 0.82 | 1.93 | 799 | 322 | 550 | 46 | 44 | 54 |
| *Streptococcus salivarius* | A0A6N3BLK3 | A0A7L6WIW1 | A0A448A7C8 | 0.75 | 0.83 | 1.51 | 916 | 382 | 533 | 48 | 45 | 56 |
| *Staphilococcus aureus* | A0A6K1MRM3 | A0A6B5I2H0 | A0A3A5M2J4 | 0.73 | 0.78 | 2.71 | 875 | 383 | 621 | 20 | 30 | 32 |
| *Bacillus subtilis* | P39695 | P39147 | P39145 | 0.76 | 0.83 | 2.58 | 1082 | 434 | 650 | 26 | 36 | 36 |
| *Lactobacillus sakei* | Q38WR1 | Q38YD4 | Q38YD5 | 0.72 | 0.79 | 2.75 | 1011 | 515 | 418 | 24 | 32 | 39 |
| *Lactococcus lactis* | Q9CER4 | Q9CGK7 | Q9CGK6 | 0.75 | 0.83 | 1.77 | 967 | 364 | 459 | 44 | 45 | 50 |
| *Staphylococcus epidermidis* | Q5HNV9 | Q5HQX8 | Q5HQX9 | 0.71 | 0.77 | 2.99 | 864 | 319 | 680 | 20 | 30 | 34 |

*: RMSD is computed between ComEC/FC/FA complexes excluding DNA and against Streptococcus pneumoniae prediction, using "matchmaker" ChimeraX module.
Iteration cutoff distance of 5 were used for pruning

**: Interacting surface area is computed using AF3 models and "measure buriedarea" ChimeraX command

***: Sequence identity with *S.pneumoniae* sequences

**Table S1: Strains and primers used in this study**

| Strain | Genotype | Source/reference |
|---|---|---|
| ***S. pneumoniae* strains** | | |
| R304 | *rpsL41, rif23, nov1* ; $Sm^R$, $Rif^R$, $Nov^R$ | 13 |
| R800 | *Non-capsulated D39 derivative, with 8,651 bp deletion in the cps locus* | 14 |
| R1331 | *recA-his; ssbB::pR424 (luc); comA::ermAM; dprA::kan* $^{184A}$ ; CmR, ErmR, KanR, SpcR | 15 |
| R1501 | *comC0* | 16 |
| R1521 | *comC0, comC-luc* ; $Ery^R$ | 5 |
| R1843 | *comC0; hexA::spc* ; $Spc^R$ | 5 |
| R2300 | *comC0, comEC::kan* $^{2a}$ ; $Kan^R$ | 8 |
| R4654 | *comC0, comC::pR414 (luc), comFAC::cat* ; $Spc^R$, $Cm^R$ | This study |
| R4857 | *comC0, ΔrecA::trim* ; $Trim^R$ | 6 |
| R4858 | *comC0, dprA-LgBit, CEP* $_{lac}$ *-dprA-SmBit* ; $Kan^R$ | 6 |
| R5000 | *comC0, comEC* $^{Y157E}$ | This study |
| R5002 | *comC0, comEC* $^{Y116E+Y157E}$ | This study |
| R5003 | *comC0, comEC* $^{F152D}$ | This study |
| R5007 | *comC0, comEC* $^{R292M}$ | This study |
| R5009 | *comC0, comEC* $^{R292K}$ | This study |
| R5011 | *comC0, comEC* $^{K297E}$ | This study |
| R5064 | *comC0, hexA::spc, comFA* $^{Y59D}$ ; $Spc^R$ | This study |
| R5099 | *comC0, Δ comFC::trim* ; $Trim^R$ | This study |
| R5114 | *comC0, comFAstop* | This study |
| R5121 | *comC0, comFAstop, CEP* $_{lac}$ *-comFC* ; $Kan^R$ | This study |
| R5153 | *comC0, comFC* $^{F95D}$ | This study |
| R5162 | *comC0, comFA -lgbit, smbit-comFC* | This study |
| R5179 | *comC0, comFA* $^{Y59D}$ *-lgbit, smbit-comFC* | This study |
| R5191 | *comC0, comFC* $^{K83D}$ | This study |
| R5194 | *comC0, comFA -lgbit* | This study |
| R5238 | *comC0, comFC* $^{R156D}$ | This study |
| R5240 | *comC0, comFA-lgbit, smbit-comFC* $^{F95D}$ | This study |
| R5251 | *comC0, comFA* $^{Y59D}$ *, comFC* $^{K83D}$ | This study |

| | | |
|---|---|---|
| R5252 | *comC0, comFA-lgbit, smbit-comFC, comEC::kan* $^{2a}$, Kan$^{R}$ | This study |
| R5255 | *comC0, comEC* $^{alfa349}$ | This study |
| R5257 | *comC0, comFA-lgbit, smbit-comFC* $^{K83D}$ | This study |
| R5270 | *comC0, comFC* $^{K83D\text{-}R156D}$ | This study |
| R5278 | *comC0, comFA* $^{W273A}$ | This study |
| R5279 | *comC0, comFC* $^{F16D}$ | This study |
| R5280 | *comC0, comFC* $^{\Delta 12\text{-}26}$ | This study |
| R5281 | *comC0, comEC* $^{R261A}$ | This study |
| R5282 | *comC0, comEC* $^{E267K}$ | This study |
| R5283 | *comC0, comEC* $^{D457K}$ | This study |
| R5304 | *comC0, comEC* $^{Y116E}$ | This study |
| R5305 | *comC0, comFA* $^{Y59D}$, *comFC* $^{F95D}$ | This study |
| R5307 | *comC0, comFA* $^{W273A}$, *comFC* $^{F16D}$ | This study |
| R5309 | *comC0, comFA* $^{Y59D}$, *comFC* $^{F16D}$ | This study |
| R5355 | *comC0, comEC* $^{Y116E\text{-}Y157E\text{-}alfa349}$ | This study |
| R5356 | *comC0, comEC* $^{F152D\text{-}alfa349}$ | This study |
| R5357 | *comC0, comEC* $^{R292M\text{-}alfa349}$ | This study |
| R5358 | *comC0, comEC* $^{R292K\text{-}alfa349}$ | This study |
| R5359 | *comC0, comEC* $^{K297E\text{-}alfa349}$ | This study |
| R5360 | *comC0, comEC* $^{E267K\text{-}alfa349}$ | This study |
| R5361 | *comC0, comEC* $^{D457K\text{-}alfa349}$ | This study |
| R5362 | *comC0, comEC* $^{R261A\text{-}alfa349}$ | This study |
| R5363 | *comC0, hexA::spc, comEC* $^{R261A}$ ; Spc$^{R}$ | This study |
| R5366 | *comC0, comEC* $^{D457K}$, *comFC* $^{F16D}$ | This study |
| R5367 | *comC0, comFA* $^{W273D}$ | This study |
| R5368 | *comC0, hexA::spc, comFA* $^{W273D}$, *comEC* $^{R261A}$ | This study |
| R5372 | *comC0, comFC* $^{F16D+K83D}$ | This study |
| R5393 | *comC0, comEC* $^{R261A}$, *comFC* $^{F16D}$ | This study |
| R5395 | *comC0, comFA* $^{Y59D+W273A}$ | This study |
| R5431 | *comC0, comEC* $^{alfa349\text{-}D457K}$, *comFC* $^{F16D}$ | This study |
| R5432 | *comC0, comEC* $^{alfa349\text{-}R261A}$, *comFC* $^{F16D}$ | This study |
| R5433 | *comC0, comEC* $^{alfa349\text{-}R261A}$, *comFA* $^{W273D}$ | This study |

| | | |
|---|---|---|
| ***H. pylori* strains** | | |
| 1 | Reference strain *26695* | [17] |
| 134 | *26695*; Sm$^R$ | [18] |
| 1141 | $P_{comEC}$-*comEC-Flag, ΔcomEC*; Cm$^R$, Kan$^R$ | This study |
| 1385 | $P_{comEC}$-*comEC*$^{R239M}$-*Flag, ΔcomEC*; Cm$^R$, Kan$^R$ | This study |
| 1389 | $P_{comEC}$-*comEC*$^{R239K}$-*Flag, ΔcomEC*; Cm$^R$, Kan$^R$ | This study |
| 1387 | $P_{comEC}$-*comEC*$^{Y60E+Y82E}$-*Flag, ΔcomEC*; Cm$^R$, Kan$^R$ | This study |
| 1435 | $P_{ureA}$-*comEC-2ALFA, ΔcomEC*; Cm$^R$, Kan$^R$ | This study |
| 1439 | $P_{ureA}$-*comEC*$^{R239M}$-*2ALFA, ΔcomEC*; Cm$^R$, Kan$^R$ | This study |
| 1454 | $P_{ureA}$-*comEC*$^{R239K}$-*2ALFA, ΔcomEC*; Cm$^R$, Kan$^R$ | This study |
| 1456 | $P_{ureA}$-*comEC*$^{Y60E+Y82E}$-*2ALFA, ΔcomEC*; Cm$^R$, Kan$^R$ | This study |
| ***Escherichia coli* strains (DH5α)** | | |
| 1862 | pJET1.2-*p_comEC-comEC*$_{Hp}^{R239M}$-*Flag* | This study |
| 1863 | pJET1.2-*p_comEC-comEC*$_{Hp}^{R239K}$-*Flag* | This study |
| 1864 | pJET1.2-*p_comEC-comEC*$_{Hp}^{Y60E+Y82E}$-*Flag* | This study |
| 1868 | pJET1.2-*p_ureA-comEC*$_{Hp}$-*2ALFA* | This study |
| 1872 | pJET1.2-*p_ureA-comEC*$_{Hp}^{R239M}$-*2ALFA* | This study |
| 1873 | pJET1.2-*p_ureA-comEC*$_{Hp}^{R239K}$-*2ALFA* | This study |
| 1874 | pJET1.2-*p_ureA-comEC*$_{Hp}^{Y60E-Y82E}$-*2ALFA* | This study |

| Primer | Sequence (5'-3') | Source/reference | Use |
|---|---|---|---|
| ***S. pneumoniae* primers** | | | |
| MB117 | AATCTCCGCTGTAGGTCACTTTCTT | [15] | Amplification of 3,434 kb fragment containing *rpsL* gene F |
| MB120 | TTGGATTGGGTGTGCATTTGC | [15] | Amplification of 3,434 kb fragment containing *rpsL* gene R |
| MB593 | TAGAATTTCTTCAAAAAGTCTATAACCTGTAAC | This study | Amplification of *smBit* R |
| OCN121 | GGTCAATGGTGATAGTCTGT | This study | Amplification of *comEC* for screening and sequencing F |
| OCN470 | CCAGCAGTAGCGACAATCTCAGCT | This study | Forward primer for construction of *comEC* mutants |
| OCN471 | CCTCGGACTGGAGTTTATA**TTC**GACTTGAAAATGCGTCCATCAGCCT | This study | Reverse primer for *comEC*$^{Y116E}$ mutagenesis |
| OCN472 | GCTGATGGACTCATTTTTCAGGTC**GAA**TATAA1ACTCCAGTCCGAGGAGGA | This study | Forward primer for *comEC*$^{Y116E}$ mutagenesis |
| OCN473 | GTGACTGTCACAGCAGATACACTGGA | This study | Amplification of 3' *comEC* for generation of comEC mutants R |
| OCN474 | GAGTCTTCAGATAGGCTTG**TTC**ATTAAAGCCACCAAAATTTCTCTGCCCT | This study | Reverse primer for *comEC*$^{Y157E}$ mutagenesis |

| | | | |
|---|---|---|---|
| OCN475 | CAGAGAAATTTTGGTGGCTTTAT**GAA**CAGCCTATCTGAAGACTCAGGA | This study | Forward primer for *comEC*$^{Y157E}$ mutagenesis |
| OCN476 | GGCTTGGTAATTAAAGCCACC**ATC**ATTTCTCTGCCCTTCTGGCT | This study | Reverse primer for *comEC*$^{F152D}$ mutagenesis |
| OCN477 | GCCAGAAGGGCAGAGAAAT**GAT**GGTGGCTTTAATTACCAAGCCT | This study | Forward primer for *comEC*$^{F152D}$ mutagenesis |
| OCN480 | GTAGCTTTTGCAAGAGACT**CAT**AATAACCGATGCTGAAAATCCAGTTAGT | This study | Reverse primer for *comEC*$^{R292M}$ mutagenesis |
| OCN481 | CTGGATTTTCAGCATCGGTTATT**ATG**AGTCTCTTGCAAAAGCTACTGGCT | This study | Forward primer for *comEC*$^{R292M}$ mutagenesis |
| OCN482 | ACCCCATGTTGAGCCAGTAG**CTC**TTGCAAGAGACTGCGAATAACCGA | This study | Reverse primer for *comEC*$^{K297E}$ mutagenesis |
| OCN483 | CGGTTATTCGCAGTCTCTTGCAA**GAG**CTACTGGCTCAACATGGGGT | This study | Forward primer for *comEC*$^{K297E}$ mutagenesis |
| OCN484 | GCCTCGGACAAGGGTAATACACGT | This study | Amplification of 5' comEC for generation of comEC mutants F |
| OCN485 | CGGATGGGAAGACTGATATAGTGA | This study | Reverse primer for construction of *comEC* mutants |
| oALS30 | GTAGCTTTTGCAAGAGACT**TTT**AATAACCGATGCTGAAAATCCAGTTAGT | This study | Reverse primer for *comEC*$^{R292K}$ mutagenesis |
| oALS31 | CTGGATTTTCAGCATCGGTTATT**AAA**AGTCTCTTGCAAAAGCTACTGGCT | This study | Forward primer for *comEC*$^{R292K}$ mutagenesis |
| OMB12 | ACAAGACGAGTTTCCAACTCTCTCTCC | This study | Amplification of 3' *comEC* for generation of comEC mutants R |
| DJ21 | AAGTAAGACCATCCCCATGCTC | This study | Amplification of 3' *comEC* for generation of comEC mutants R |
| CJ344 | TTCATAGACCAGCCTCCTTATTCATCA | This study | Sequencing of comFA mutants R |
| CJ359 | GAGATTGTGATTTTCTAGTACCCCA | This study | Amplification of 5' *comFAC* for sequencing/screening F |
| CJ362 | ATCGCTCTATGACTGACTCCAACTCC | This study | Sequencing of comFAC mutants R |
| CJ364 | TCCAACTCTTGTTCTGGTTGGAAGT | This study | Amplification of 3' *comFAC* for sequencing/screening R |
| CJ868 | AGTAGTTGTGAGAATGTCAACGAAA | This study | Amplification of 5' *comFAC* for generation of comFAC mutants F |
| CJ873 | TATAAAATTAGCTACAAAACTATTG | This study | Amplification of 3' *comFAC* for generation of comFAC mutants R |
| CJ907 | CATTCTATTTTTGCAAATTTTTTCCAAGCTTATCGATACCGTCGACCTCG | This study | Creation of *ΔcomFC::trim* R |
| CJ908 | CGAGGTCGACGGTATCGATAAGCTTGGAAAAAATTTGCAAAAATAGAATG | This study | Creation of *ΔcomFC::trim* F |
| CJ910 | TCATCAGCAAGCACTCTCGACAAT**C**GTAAGCACCGATGGGCAAATAC | This study | Creation of *comFA*$^{Y59D}$ R |
| CJ911 | GTATTTGCCCATCGGTGCTTAC**G**ATTGTCGAGAGTGCTTGCTGATGA | This study | Creation of *comFA*$^{Y59D}$ F |
| CJ926 | AACTAGTGGATCCCCCGGGCTGCAGTCATAGACCAGCCTCCTTATTCATC | This study | Creation of *ΔcomFC::trim* R |
| CJ927 | GATGAATAAGGAGGCTGGTCTATGACTGCAGCCCGGGGGATCCACTAGTT | This study | Creation of *ΔcomFC::trim* F |
| CJ942 | ATAAACGACCGAGATAATCTAA**CTACTA**TTTCATACTTCTTTATTCGTAA | This study | Creation of *comFA*$^{STOP}$ R |
| CJ943 | TTACGAATAAAGAAGTATGAAA**TAGTAG**TTAGATTATCTCGGTCGTTTAT | This study | Creation of *comFA*$^{STOP}$ F |
| MD1 | TGAAGCGAAAACTTTTCTTAACAG**ATC**GTCTCCATCAAACTTATACCGACT | This study | Creation of *comFC*$^{F95D}$ R |
| MD2 | AGTCGGTATAAGTTTGATGGAGAC**GAT**CTGTTAAGAAAAGTTTTCGCTTCA | This study | Creation of *comFC*$^{F95D}$ F |
| MD3 | ATCTGAATTTCCTTAATCGCCTTCT | This study | Generation of *comFA-LgBit, SmBit-comFC* R |
| MD4 | AGAAGGCGATTAAGGAAATTCAGAT | This study | Generation of *comFA-LgBit, SmBit-comFC* F |
| MD5 | CTAAAAGTTAAAACAGTCTTCATAG | This study | Generation of *comFA-LgBit, SmBit-comFC* R |
| MD6 | CTATGAAGACTGTTTTAACTTTTAG | This study | Generation of *comFA-LgBit, SmBit-comFC* F |

| | | | |
|---|---|---|---|
| MD39 | GAAGTATGAAA**TAGTAG** | This study | Amplification of *comFA*$^{STOP}$ F |
| MD40 | TCTGCCCACATAACAAGCACTTCATAGACCAGCCTCCTCTAACTGTTTAT | This study | Creation of *comFA-LgBit* R |
| MD41 | ATAAACAGTTAGAGGAGGCTGGTCTATGAAGTGCTTGTTATGTGGGCAGA | This study | Creation of *comFA-LgBit* F |
| MD46 | ACTTATACCGACTGAAAAAATC**ATC**CATAGCTTGATTGTAAGTAAAA | This study | Creation of *comFC*$^{K83D}$ R |
| MD47 | TTTTACTTACAATCAAGCTATG**GAT**GATTTTTTCAGTCGGTATAAGT | This study | Creation of *comFC*$^{K83D}$ F |
| MD48 | TCAATCTACGTCTAAGCTCTTCCTC | This study | Amplification of *comEC*$^{alfa349}$ R |
| MD58 | AACGATTTTTAGAAGAACTGGC**ATC**CTCTTCTCTTTTCTCTAATAAA | This study | Creation of *comFC*$^{R156D}$ R |
| MD59 | TTTATTAGAGAAAAGAGAAGAG**GAT**GCCAGTTCTTCTAAAAATCGTT | This study | Creation of *comFC*$^{R156D}$ F |
| MD68 | ATAAGTCAGCCATTTCAACTTTTCTTGGGTCAAGCCCAATCGCAAGAGAA | This study | Generation of *comEC*$^{alfa349}$ R |
| MD69 | TTCTCTTGCGATTGGGCTTGACCCAAGAAAAGTTGAAATGGCTGACTTAT | This study | Generation of *comEC*$^{alfa349}$ F |
| MD70 | TTGGGTTGTCCAAAGACCAGAGGTCTACTTGTCACCTGTGACACCAAGCG | This study | Generation of *comEC*$^{alfa349}$ R |
| MD71 | CGCTTGGTGTCACAGGTGACAAGTAGACCTCTGGTCTTTGGACAACCCAA | This study | Generation of *comEC*$^{alfa349}$ F |
| MD78 | AGTAGCGATTAAAATCCGATAA**AGC**AATTGGTTTTGGAATAATCAAC | This study | Creation of *comFA*$^{W273A}$ R |
| MD79 | GTTGATTATTCCAAAACCAATT**GCT**TTATCGGATTTTAATCGCTACT | This study | Creation of *comFA*$^{W273A}$ F |
| MD80 | TCCTCAGAAGTAAGAGACTACT**ATC**AGTTAAAACAGTCTTCATAGTC | This study | Creation of *comFC*$^{F16D}$ R |
| MD81 | GACTATGAAGACTGTTTTAACT**GAT**AGTAGTCTCTTACTTCTGAGGA | This study | Creation of *comFC*$^{F16D}$ F |
| MD82 | ACTTTTCTTGGGTCAAGCCCAA**AGC**CAAGAGAAGTTTCTTAAATCCA | This study | Creation of *comEC*$^{R261A}$ R |
| MD83 | TGGATTTAAGAAACTTCTCTTG**GCT**TTGGGCTTGACCCAAGAAAAGT | This study | Creation of *comEC*$^{R261A}$ F |
| MD84 | GATAAGTCAGCCATTTCAACTT**CTT**TTGGGTCAAGCCCAATCGCAAG | This study | Creation of *comEC*$^{E267K}$ R |
| MD85 | CTTGCGATTGGGCTTGACCCAA**AAG**AAGTTGAAATGGCTGACTTATC | This study | Creation of *comEC*$^{E267K}$ F |
| MD86 | AGTCTGAACAAAGACAAGA**ACCACCACTACC**CTTCATAGTCTGCCCACAT | This study | Creation of *comFC*$^{\Delta 12\text{-}26}$ R |
| MD87 | ATGTGGGCAGACTATGAAG**GGTAGTGGTGGT**TCTTGTCTTTGTTCAGACT | This study | Creation of *comFC*$^{\Delta 12\text{-}26}$ F |
| MD88 | GCTTTTTAATGTTTTTCCTCAA**TTT**ATAGACCAAAGCCAAAGAAATT | This study | Creation of *comEC*$^{D457K}$ R |
| MD89 | AATTTCTTTGGCTTTGGTCTAT**AAA**TTGAGGAAAAACATTAAAAAGC | This study | Creation of *comEC*$^{D457K}$ F |
| MD102 | AGTAGCGATTAAAATCCGATAA**ATC**AATTGGTTTTGGAATAATCAAC | This study | Creation of *comFA*$^{W273D}$ R |
| MD103 | GTTGATTATTCCAAAACCAATT**GAT**TTATCGGATTTTAATCGCTACT | This study | Creation of *comFA*$^{W273D}$ F |
| OEC060 | CCAGATTAGGAACTATCCGTGAGT | This study | Creation of *ΔcomFAC* |
| OEC061 | ATGTGACCAGTCAGCCTCGCCATACTTCTTTATTCGTAAAAACTAGC | This study | Creation of *ΔcomFAC* |
| OEC062 | GCTAGTTTTTACGAATAAAGAAGTATGGCGAGGCTGACTGGTCACAT | This study | Creation of *ΔcomFAC* |
| OEC063 | TACATCCTTAGCACCGGCTTCGAAGGATATGGATCTGGAGCTGT | This study | Creation of *ΔcomFAC* |
| OEC064 | ACAGCTCCAGATCCATATCCTTCGAAGCCGGTGCTAAGGATGTA | This study | Creation of *ΔcomFAC* |
| CJ356 | ATACAAAACACCGGAAGAAGCATCC | This study | Creation of *ΔcomFAC* |

***H. pylori* primers**

| | | | |
|---|---|---|---|
| OC178 | CAAATCTTGTTGCAA**GAG**CCTAAAACTAAAGATCAAAAAACCTATTTTGT | This study | *comEC* $^{Y60E}$ mutagenesis F |
| OC179 | ATCTTTAGTTTTAGG**CTC**TTGCAACAAGATTTGAGCGCTCAAACTTGTAG | This study | *comEC* $^{Y60E}$ mutagenesis R |
| OC174 | AAAAACATGATCTTT**GAG**ACCACCATTAAAGAGCCTTTAAAAAAACCTCCA | This study | *comEC* $^{Y82E}$ mutagenesis F |
| OC175 | TAATGGTGGT**CTC**AAAGATCATGTTTTTTGATTGGAGCTTTAAAAC | This study | *comEC* $^{Y82E}$ mutagenesis R |
| OC176 | CCCTCTTTTTTC**ATG**GCGTTTTTAATGGGCTTATTAGGGTTTTTGGCATG | This study | *comEC* $^{R239M}$ mutagenesis F |
| OC177 | CATTAAAAACGC**CAT**GAAAAAAGAGGGTAAAAAATCTAGTAGCAACAAAT | This study | *comEC* $^{R239M}$ mutagenesis R |
| OC151 | CCCTCTTTTTTC**AAA**GCGTTTTTAATGGGCTTATTAGGGTTTTTGGCATG | This study | *comEC* $^{R239K}$ mutagenesis F |
| OC152 | CATTAAAAACGC**TTT**GAAAAAAGAGGGTAAAAAATCTAGTAGCAACAAAT | This study | *comEC* $^{R239K}$ mutagenesis R |
| OC153 | AAGGAAAAACACTTTAAGAATAGGAGAATAAGTTGAAAGACAAAACTTTTCAGGGG | This study | Insertion 2ALFA Tag in Cter on *comEC* F (PCR1) |
| OC154 | AGCTCTTCCTCCAATCGCGAAGGCTCAGTTAAACGGCGCCGGAGTTCCTCTTCAAGCCTTGACGGACCT ACAATTAAGCTAGGCATTATA | This study | Insertion 2ALFA Tag in Cter on *comEC* R (PCR1) |
| OC156 | CCCCTGAAAAGTTTTGTCTTTCAACTTATTCTCCTATTCTTAAAGTGTTTTTCCTT | This study | Insertion 2ALFA Tag in C-ter on *comEC* R (PCR2) |
| OC111 | AGAGCTTCGCCGTCGGCTAACAGAAtagGCCATATTGTGTTGAAACACCGCCC | This study | Insertion 2ALFA Tag in Cter on *comEC* F (PCR2) |
| AM10 | TTATTAAACGAGCGCCATTCTTGC | This study | Insertion check in *rdxA* locus F |
| AM11 | CCGGATAGAGATTTTGCATGTA | This study | Insertion check in *rdxA* locus R |
| AM1 | GGTCGCGTAGCTCAGTTGGT | This study | Insertion check in *ureA* locus F |
| AM13 | CCTCAATAGGGGTATGC | This study | Insertion check in *ureA* locus R |
| OP137 | CGGTTTTTCGTCCTTGGGTTATC | This study | Insertion check in *comEC* locus R |
| OP138 | GTGTGTGGGGTGTTTTTAAGCC | This study | Insertion check in *comEC* locus F |
