## Supplementary figures and images for "A tripartite protein complex promotes DNA transport during natural transformation in firmicutes"

### Dewailly Fauconnet 2025 - Supplementary Figures

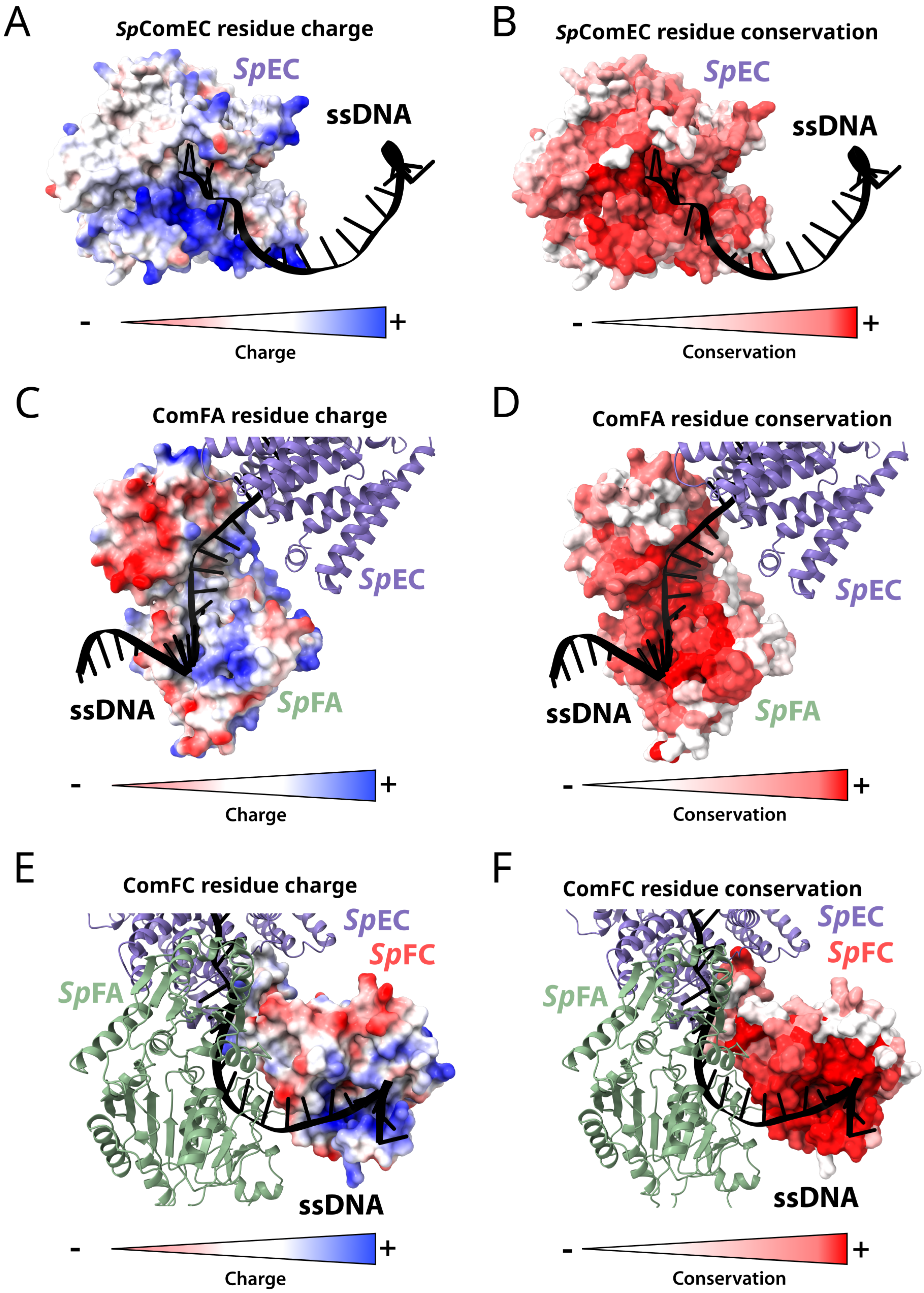

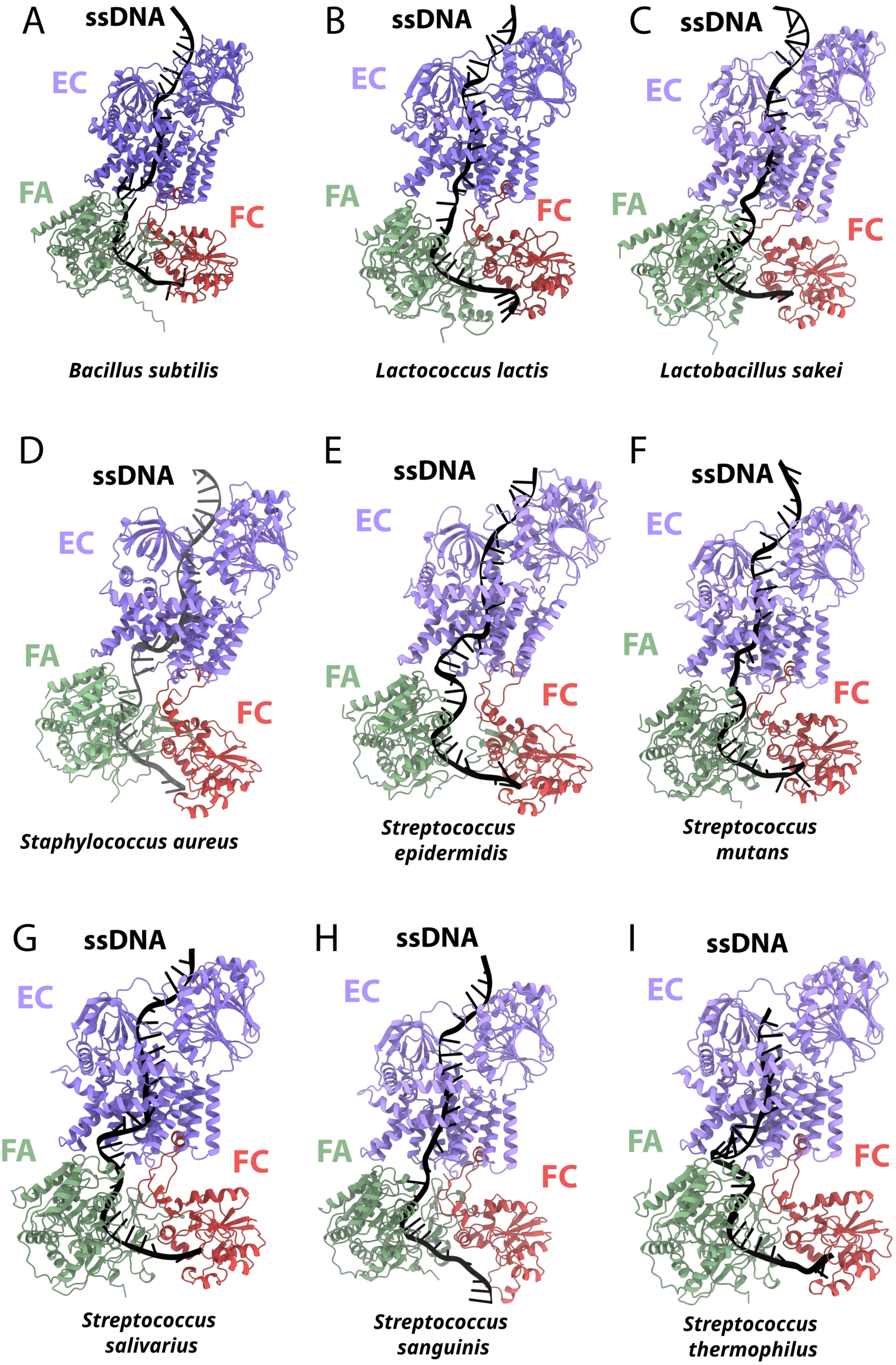

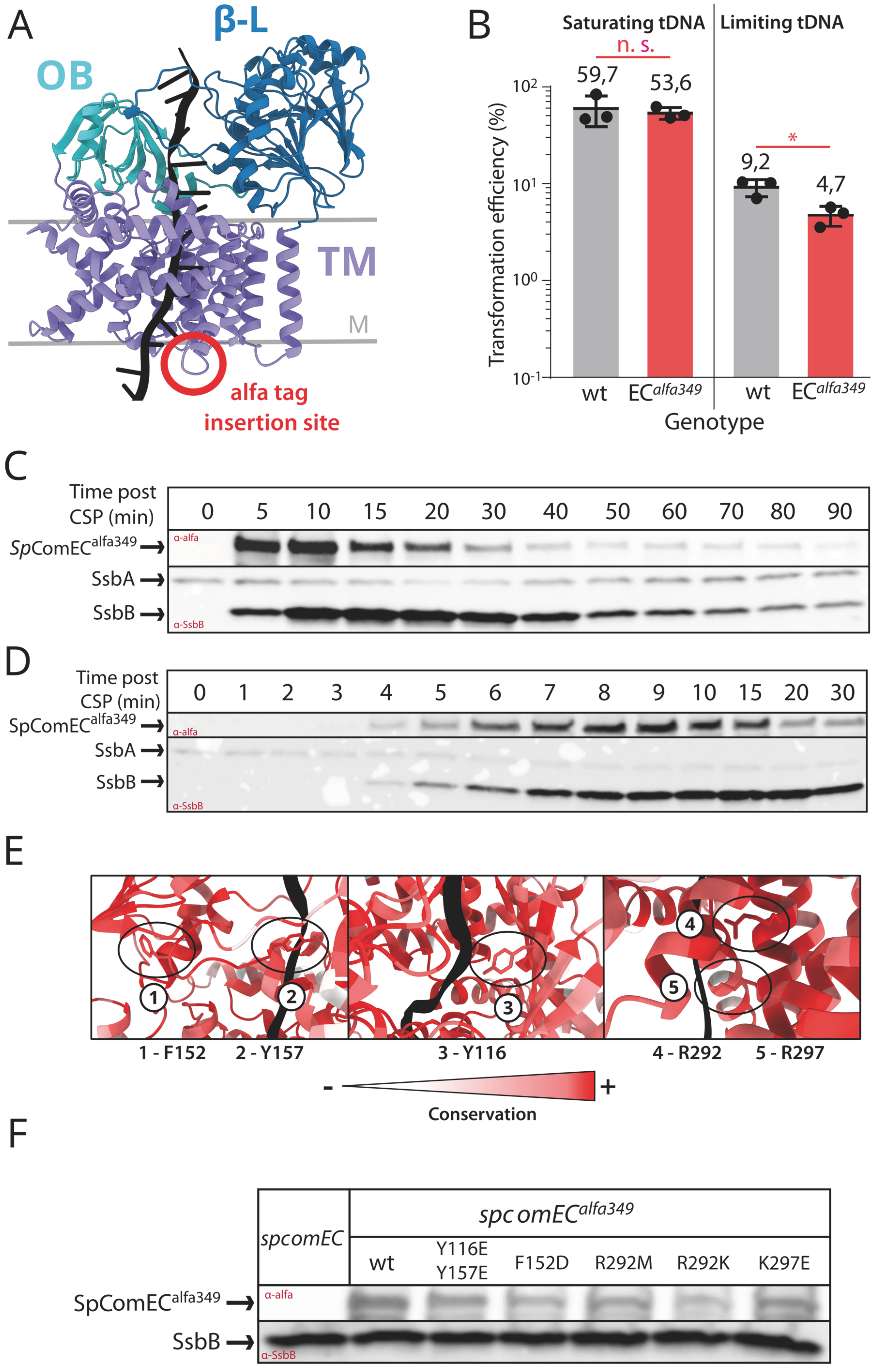

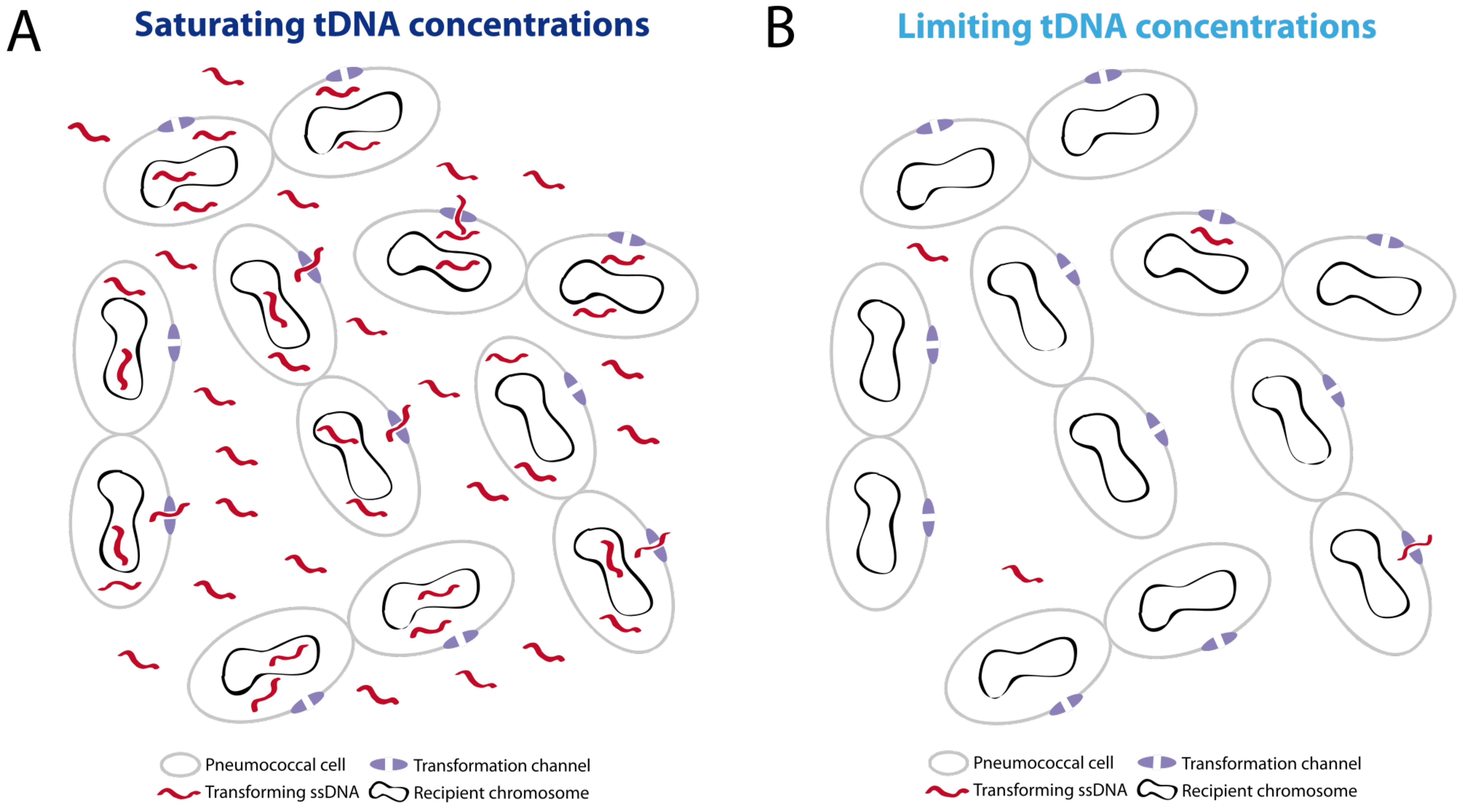

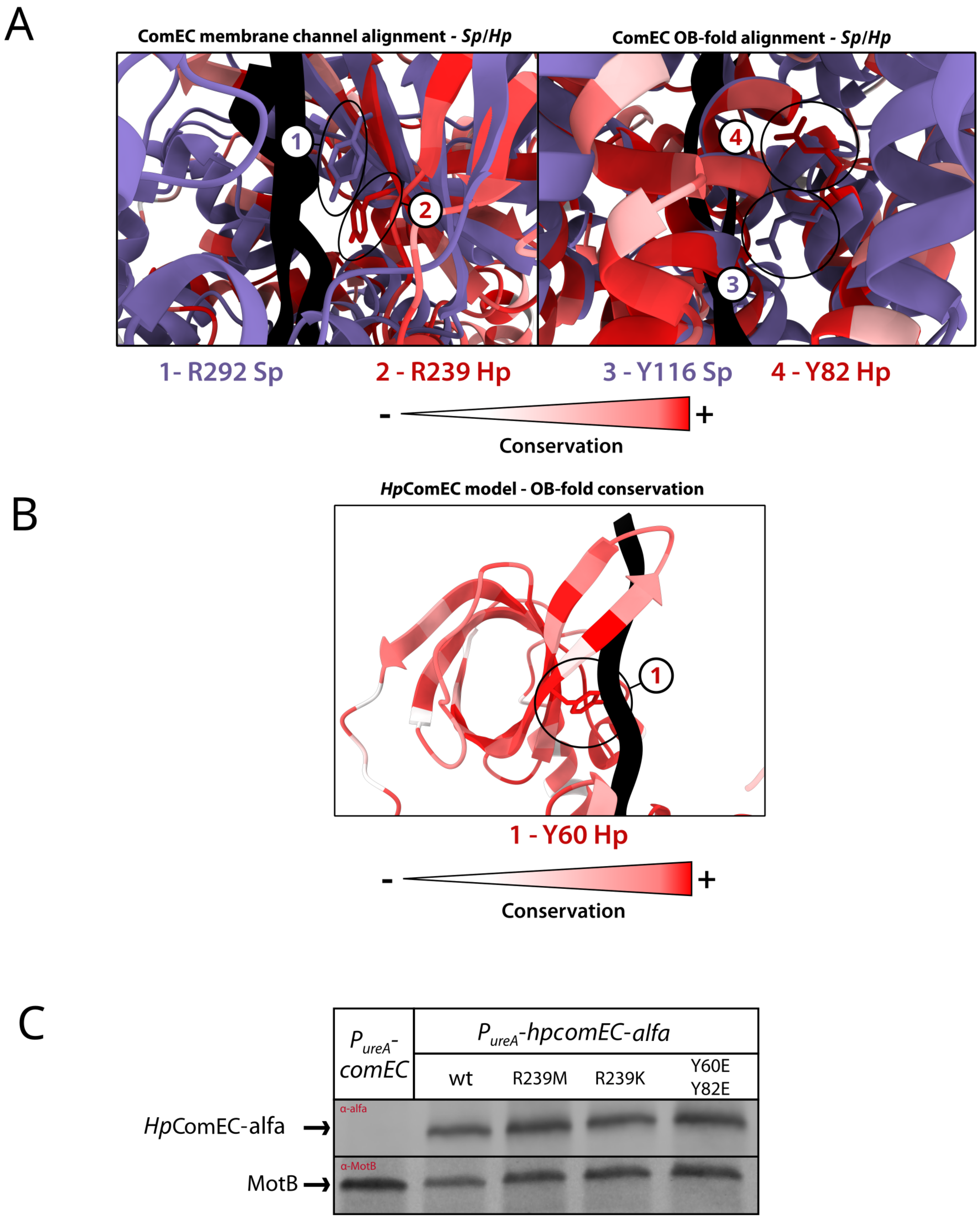

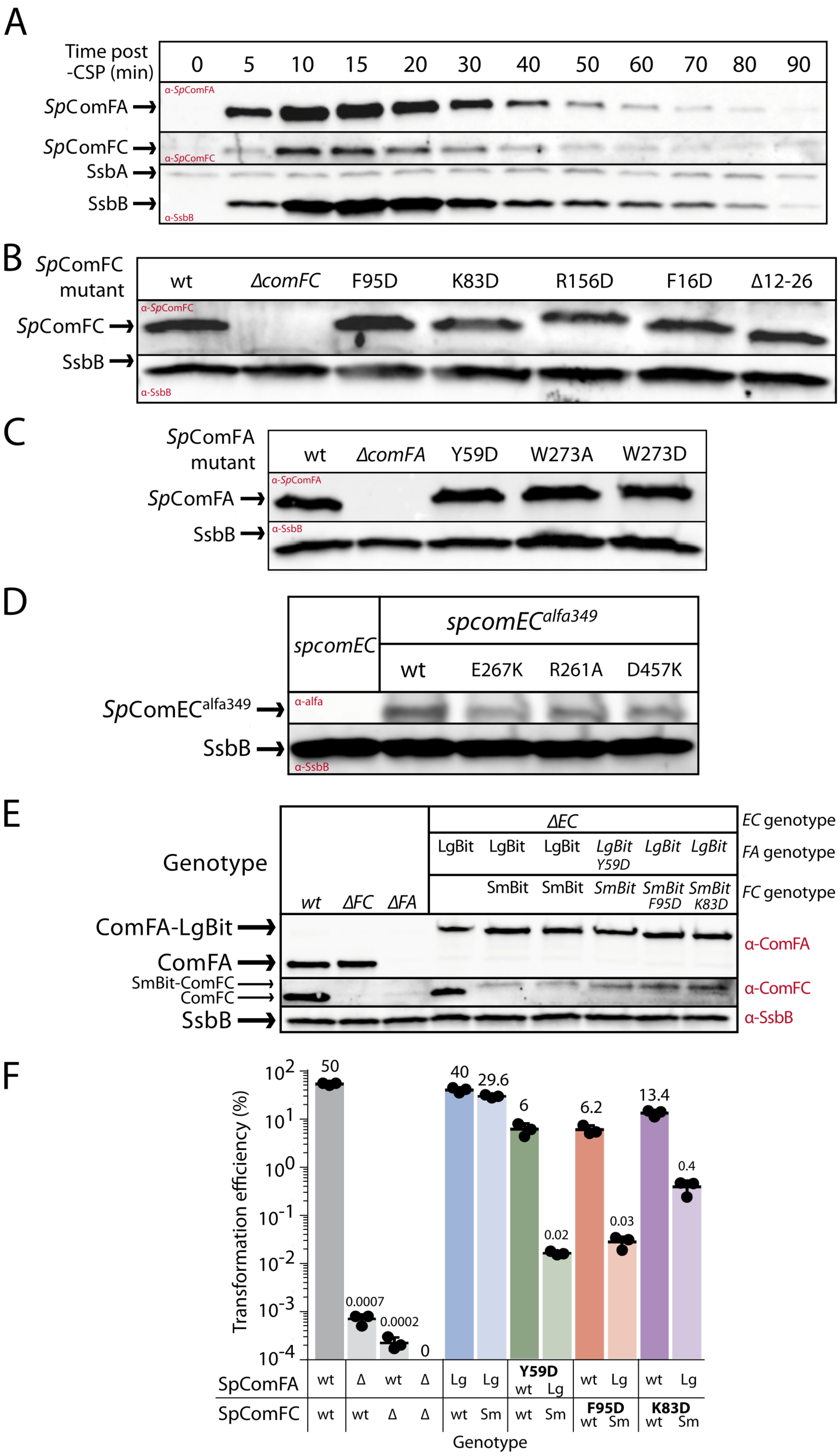

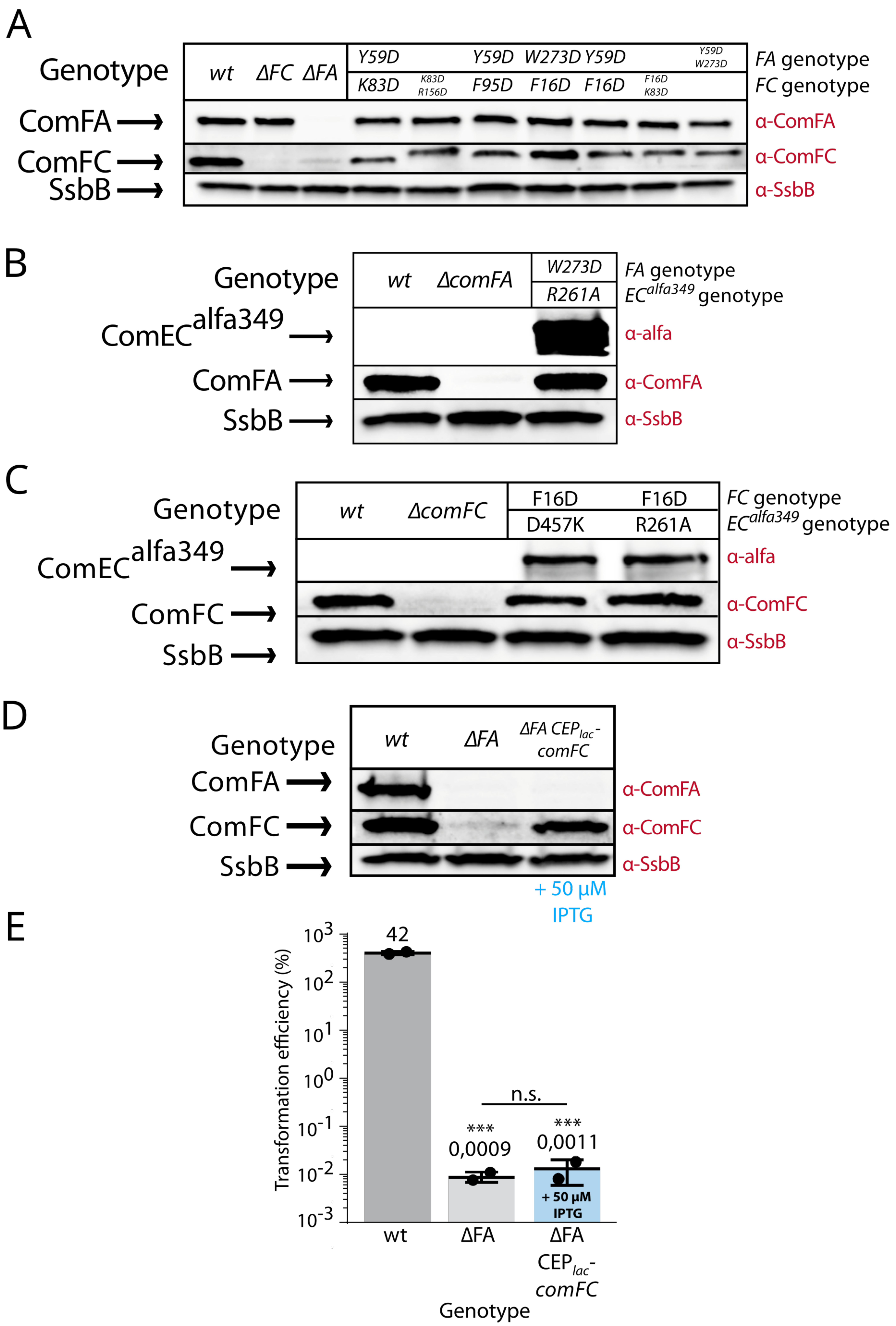

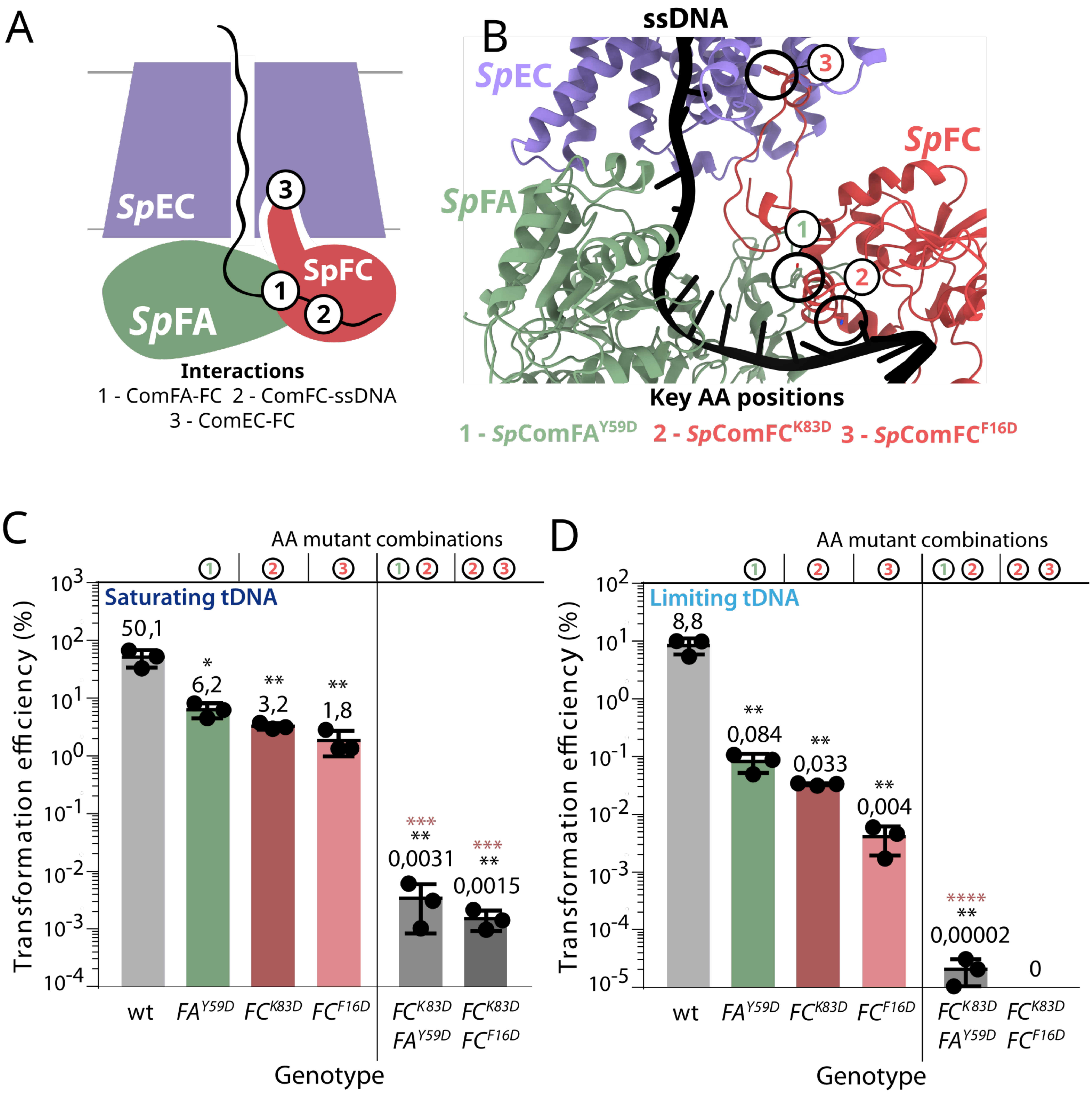
