## Supplementary material for "A tripartite protein complex promotes DNA transport during natural transformation in firmicutes": Dewailly Fauconnet 2025 - Supplementary Information

This PDF file includes:

- Supplementary Methods
- Supplementary Results and Discussion
- Supplementary Figures S1-S8
- Tables S1-S3

### Supplementary Methods

**Construction of *S. pneumoniae* mutant strains.**

Here we describe how the new mutant strains used in this study were generated. Previously published constructs and mutants were simply transferred from published strains by transformation with appropriate selection. To obtain certain mutant strains, it was necessary to delete the hexA gene, thereby suppressing the Hex generalised mismatch repair system[1,2]. This has been shown to slightly affect the transformation efficiency of the *rpsL41* point mutation[3,4], but since this was a minor effect and the transformation deficits were large in these strains, this was not taken into account. Strains generated and primers used can be found in Table S3.

To generate the R4654 strain (*comC0, comC-luc, comFAC::cat*), three PCR fragments were generated using the following primer pairs and matrixes: OCE060-OEC061 on R1501 gDNA, OCE062-OEC063 on R1331 gDNA and OEC064-CJ356 on R1501 gDNA. These represented the upstream sequence of the *spcomFAC* operon, a chloramphenicol resistance cassette and the downstream sequence of the *spcomFAC* operon respectively. A SOE PCR fragment generated using these three fragments was transformed into R1521[5] and resulting transformants selected using chloramphenicol.

The R5000 strain (*comC0*, $SpcomEC^{Y157E}$) was generated by amplifying two adjacent DNA fragments by PCR on the R1501 strain in the *SpcomEC* gene using primer pairs OCN470-OCN474 and OCN475-OCN485 respectively. Splicing overlap extension (SOE) PCR on these two fragments with the OCN470-OCN485 primer pair generated a DNA fragment with the $SpcomEC^{Y157E}$ mutation. This DNA fragment was transformed without selection into R1501 with a 3 h 30 min phenotypic expression phase in liquid culture to introduce the $SpcomEC^{Y157E}$ mutation and positive clones were determined by PCR amplification with the OCN121-OCN473 primer pair and sequencing with the OCN121 primer.

The R5002 strain (*comC0, SpcomEC*$^{Y116E+Y157E}$) was generated as described for strain R5000, except that primer pairs OCN470-OCN471 and OCN472-OCN485 were used for initial PCRs on strain R5000.

The R5003 strain (*comC0, SpcomEC*$^{F152D}$) was generated as described for strain R5000, except that primer pairs OCN470-OCN476 and OCN477-OCN485 were used for initial PCRs.

The R5007 strain (*comC0, SpcomEC*$^{R292M}$) was generated as described for strain R5000, except that primer pairs OCN470-OCN480 and OCN481-OCN485 were used for initial PCRs.

The R5009 strain (*comC0, SpcomEC*$^{R292K}$) was generated as described for strain R5000, except that primer pairs OCN470-oALS30 and oALS31-OCN485 were used for initial PCRs.

The R5011 strain (*comC0, SpcomEC*$^{K297E}$) was generated as described for strain R5000, except that primer pairs OCN470-OCN482 and OCN483-OCN485 were used for initial PCRs.

The R5064 strain (*comC0, hexA::spc, comFA*$^{Y59D}$) was generated by amplifying two adjacent DNA fragments by PCR on the R1501 strain in the *comFA* gene using primer pairs CJ868-CJ910 and CJ911-CJ873 respectively. Splicing overlap extension (SOE) PCR on these two fragments with the CJ868-CJ873 primer pair generated a DNA fragment with the *comFA*$^{Y59D}$ mutation. This DNA fragment was transformed without selection into R1843 with a 3 h 30 min phenotypic expression phase in liquid culture to introduce the *comFA*$^{Y59D}$ mutation and positive clones were determined by PCR amplification with the CJ359-CJ364 primer pair and sequencing with the CJ359 primer.

To generate the R5099 strain (*comC0, ∆comFC::trim*), a *∆comFC::trim* DNA fragment was created by initial amplification of the regions upstream and downstream of the comFC gene using primer pairs CJ868-CJ926 and CJ908-CJ873 and the R1501 strain as template. The trimethoprim resistance cassette was amplified using the primer pair CJ927-CJ907 and the R4857 strain[6] as template. SOE PCR on these three fragments with the primer pair CJ868-CJ873 generated a DNA fragment with the *comFC* gene replaced with the trimethoprim resistance cassette, which was transformed into R1501 and transformants were selected with trimethoprim. The absence of *comFC* expression in this strain was validated by Western blot (Figure S5B).

To generate the R5111 strain (*comC0, CEP*$_{lac}$*-comFC*), a *CEP*$_{lac}$*-comFC* DNA fragment was created by initial amplification of the regions upstream and downstream of the CEP platform using primer pairs CJ574-CJ934 and CJ937-CJ575 and the R4848[6] strain as template. The *comFC* gene was amplified using the primer pair CJ935-CJ936 and the R1501 strain as template. SOE PCR on these three fragments with the primer pair CJ574-CJ575 generated a DNA fragment with the *comFC* gene controlled by the $P_{lac}$ promoter, which was transformed into R1501 and transformants were selected with kanamycin.

To inactivate *comFA* while affecting *comFC* expression as little as possible, we replaced codons 3 and 4 of the ComFA protein with stop codons. The R5114 strain (*comC0, comFA*$^{stop}$) was generated by amplifying two adjacent DNA fragments by PCR on the R1501 strain in the *comFA* gene using primer pairs CJ868-CJ942 and CJ943-CJ873 respectively. The six base mutations required to alter V3 and N4 to two stop codons are present in both primers CJ942 and CJ943. SOE PCR on these two fragments with the CJ868-CJ873 primer pair generated a DNA fragment with the *comFA*$^{stop}$ mutation. This DNA fragment was transformed without selection into R1501 with a 3 h 30 min phenotypic expression phase in liquid culture to introduce the *comFA*$^{stop}$ mutation and positive clones were determined by PCR amplification with the MD39-CJ364 primer pair and sequencing with the CJ359 primer. The absence of *comFA* expression in this strain was validated by Western blot (Figure S5C).

To generate the R5121 strain (*comC0, CEP*$_{lac}$*-comFC, comFA*$^{stop}$), R5111 was transformed with a DNA fragment containing the *comFA*$^{stop}$ mutation (generated as previously described for strain R5114) without selection and positive clones were determined by PCR amplification with the MD39-CJ364 primer pair and sequencing with the CJ359 primer. The absence of *comFA* expression and the *comFC* expression in this strain with 50 µM were validated by Western blot (Figure S7D).

The R5153 strain (*comC0*, $comFC^{F95D}$) was generated as described for strain R5064, except that primer pairs CJ868-MD1 and MD2-CJ873 were used for initial PCRs, the CJ868-CJ873 primer pair used for SOE PCR, and transformants identified by PCR with the CJ359-CJ364 primer pair and sequencing with the CJ364 primer.

To generate strain R5162 (*comC0, comFA-lgbit, smbit-comFC*), a *comFA-lgbit, smbit-comFC* DNA fragment was created by initial amplification of the regions upstream and downstream of the *comFAC* genes using primer pairs CJ868-MD3 and MD6-CJ873 and R1501 gDNA as a template. A DNA fragment containing the *lgbit* and *smbit* tag sequences in frame with *comFA* and *comFC* respectively, separated by appropriate linkers was synthesized based on previously-published sequences optimized for the pneumococcus[6,7] using gBlocks (Integrated DNA technologies). The *comFA-lgbit, smbit-comFC* operon was amplified using the primer pair MD4-MD5 and the synthetic DNA fragment *comFA-Lgbit, smbit-comFC* as template. SOE PCR on these three fragments with the primer pair CJ868-CJ873 generated a *comFA-lgbit, smbit-comFC* fragment which was transformed into R1501 without selection and transformants were screened by PCR for integration using primer pair CJ359-CJ364.

Strain R5179 (*comC0*, $comFA^{Y59D}$*-lgbit, smbit-comFC*) was generated as described for strain R5064, except that primer pairs CJ868-CJ910 and CJ911-CJ873 were used for initial PCRs on strain R5162, the CJ868-CJ873 primer pair used for SOE PCR, and transformants identified by PCR with the CJ359-CJ364 primer pair and sequencing with the CJ359 primer.

The R5191 strain (*comC0*, $comFC^{K83D}$) was generated as described for strain R5064, except that primer pairs CJ868-MD46 and MD47-CJ873 were used for initial PCRs, the CJ868-CJ873 primer pair used for SOE PCR, and transformants identified by PCR with the CJ359-CJ364 primer pair and sequencing with the CJ364 primer.

The R5194 strain (*comC0, comFA-lgbit*) was generated by amplifying two adjacent DNA fragments by PCR on the R5162 strain in the comFAC genes using primer pairs CJ868-MD40 and MD41-CJ873 respectively. SOE PCR on these two fragments with the CJ868-CJ873 primer pair generated a *comFA-lgbit* fragment associated to wildtype *comFC*, which was transformed into R1501 without selection and transformants were screened by PCR for integration using primer pair CJ359-CJ364.

The R5237 strain (*comC0, comFC*$^{K11D}$) was generated as described for strain R5064, except that primer pairs CJ868-MD56 and MD57-CJ873 were used for initial PCRs, the CJ868-CJ873 primer pair used for SOE PCR, and transformants identified by PCR with the CJ359-CJ364 primer pair and sequencing with the CJ362 primer.

The R5238 strain (*comC0, comFC*$^{R156D}$) was generated as described for strain R5064, except that primer pairs CJ868-MD58 and MD59-CJ873 were used for initial PCRs, the CJ868-CJ873 primer pair used for SOE PCR, and transformants identified by PCR with the CJ359-CJ364 primer pair and sequencing with the CJ362 primer.

The R5240 strain (*comC0, comFA-lgbit-smbit-comFC*$^{F95D}$) was generated as described for strain R5064, except that primer pairs CJ868-MD1 and MD2-CJ873 were used for initial PCRs on strain R5162, the CJ868-CJ873 primer pair used for SOE PCR, and transformants identified by PCR with the CJ359-CJ364 primer pair and sequencing with the CJ364 primer.

The R5251 strain (*comC0, comFA*$^{Y59D}$*, comFC*$^{K83D}$) was generated as described for strain R5064, except that primer pairs CJ868-MD46 and MD47-CJ873 were used for initial PCRs on strain R5064, the CJ868-CJ873 primer pair used for SOE PCR, and transformants identified by PCR with the CJ359-CJ364 primer pair and sequencing with both CJ359 and CJ364 primers.

To generate strain R5252, the R5162 strain was transformed with gDNA from the R2300[8] strain and a KanR transformant was recovered.

To generate strain R5255 (*comC0, SpcomEC*$^{alfa349}$), a *SpcomEC*$^{alfa349}$ DNA fragment was created by initial amplification of the regions upstream and downstream of the *SpcomEC* gene using primer pairs OCN484-MD68 and MD71-OMB12 and R1501 gDNA as template. The alfa tag with appropriate linkers and overhang sequences for SOE PCR was synthesized based on previously-published sequence[9] optimized for the pneumococcus using gBlocks (Intergrated DNA technologies). The *SpcomEC*$^{alfa349}$ gene was amplified using the primer pair MD69-MD70 and the synthetic gene *SpcomEC*$^{alfa349}$ as a template. SOE PCR on these three fragments with the primer pair OCN484-OMB12 generated a *SpcomEC*$^{alfa349}$ fragment which was transformed into R1501 without selection and transformants were screened by PCR for integration using primer pair OCN121-MD48.

The R5257 strain (*comC0, comFA-lgbit-smbit-comFC*$^{K83D}$) was generated as described for strain R5064, except that primer pairs CJ868-MD46 and MD47-CJ873 were used for initial PCRs on strain R5162.

The R5270 strain (*comC0, comFC*$^{R156D\text{-}K83D}$) was generated as described for strain R5064, except that primer pairs CJ868-MD46 and MD47-CJ873 were used for initial PCRs on strain R5191.

The R5278 strain (*comC0, comFA*$^{W273A}$) was generated as described for strain R5064, except that primer pairs CJ868-MD78 and MD79-CJ873 were used for initial PCRs, and CJ344 primer used for sequencing transformants.

The R5279 strain (*comC0, comFC*$^{F16D}$) was generated as described for strain R5064, except that primer pairs CJ868-MD80 and MD81-CJ873 were used for initial PCRs, and CJ362 primer used for sequencing transformants.

The R5280 strain (*comC0, comFC*$^{\Delta 12\text{-}26}$) was generated as described for strain R5064, except that primer pairs CJ868-MD86 and MD87-CJ873 were used for initial PCRs, and CJ362 primer used for sequencing transformants.

The R5281 strain (*comC0, SpcomEC*$^{R261A}$) was generated as described for strain R5064, except that primer pairs OCN484-MD82 and MD83-OMB12 were used for initial PCRs, the OCN484-OMB12 primer pair used for SOE PCR, the OCN121-OCN473 primer pair used to amplify DNA from transformants and OCN121 used for sequencing transformants.

The R5282 strain (*comC0, SpcomEC*$^{E267K}$) was generated as described for strain R5281, except that primer pairs OCN484-MD84 and MD85-OMB12 were used for initial PCRs.

The R5283 strain (*comC0, SpcomEC*$^{D457K}$) was generated as described for strain R5281, except that primer pairs OCN484-MD88 and MD89-OMB12 were used for initial PCRs and OCN473 was used for transformant sequencing.

The R5304 strain (*comC0, SpcomEC*$^{Y116E}$) was generated as described for strain R5000, except that primer pairs OCN470-OCN471 and OCN472-OCN485 were used for initial PCRs.

The R5305 strain (*comC0, comFA*$^{Y59D}$*, comFC*$^{F95D}$) was generated as described for strain R5251, except that primer pairs CJ868-MD1 and MD2-CJ873 were used for initial PCRs on strain R5064.

The R5307 strain (*comC0, comFA*$^{W273A}$*, comFC*$^{F16D}$) was generated as described for strain R5064, except that primer pairs CJ868-MD78 and MD79-CJ873 were used for initial PCRs on strain R5279, R1501 was used as recipient strain, and sequencing was carried out with both CJ344 and CJ362 primers.

The R5309 strain (*comC0, hexA::spc, comFA*$^{Y59D}$*, comFC*$^{F16D}$) was generated as described for strain R5064, except that primer pairs CJ868-MD80 and MD81-CJ873 were used for initial PCRs on strain R5064, R1843 was used as recipient strain, and sequencing was carried out with both CJ359 and CJ362 primers.

The R5355 strain (*comC0, SpcomEC*$^{Y116E\text{-}Y157E\text{-}alfa349}$) was generated by amplifying two adjacent DNA fragments by PCR in the *SpcomEC* gene using primer pairs OCN470-OCN474 on R5002 strain and OCN475-OCN485 on R5255 strain respectively. Splicing overlap extension (SOE) PCR on these two fragments with the OCN470-OCN485 primer pair generated a DNA fragment with the *SpcomEC*$^{Y116E\text{-}Y157E\text{-}alfa349}$ mutation. This DNA fragment was transformed without selection into R5255 with a 3 h 30 min phenotypic expression phase in liquid culture to introduce the*SpcomEC*$^{Y116E\text{-}Y157E\text{-}alfa349}$ mutation and positive clones were determined by PCR amplification with the OCN121-OCN473 primer pair and sequencing with the OCN121 primer.

The R5356 strain (*comC0, SpcomEC*$^{F152D\text{-}alfa349}$), was generated as described for strain R5355, except that primer pairs OCN470-OCN476 and OCN477-OCN485 were used for initial PCRs on strain R5255.

The R5357 strain (*comC0, SpcomEC*$^{R292M\text{-}alfa349}$), was generated as described for strain R5355, except that primer pairs OCN470-OCN480 and OCN481-OCN485 were used for initial PCRs on strain R5255.

The R5358 strain (*comC0, SpcomEC*$^{R292K\text{-}alfa349}$), was generated as described for strain R5355, except that primer pairs OCN470-oALS30 and oALS31-OCN485 were used for initial PCRs on strain R5255.

The R5359 strain (*comC0, SpcomEC*$^{K297E\text{-}alfa349}$), was generated as described for strain R5355, except that primer pairs OCN470-OCN482 and OCN483-OCN485 were used for initial PCRs on strain R5255.

The R5360 strain (*comC0, SpcomEC*$^{E267K\text{-}alfa349}$), was generated as described for strain R5355, except that primer pairs OCN484-MD84 and MD85-OMB12 were used for initial PCRs on strain R5255.

The R5361 strain (*comC0, SpcomEC*$^{D457K\text{-}alfa349}$), was generated as described for strain R5355, except that primer pairs OCN484-MD88 and MD89-OMB12 were used for initial PCRs on strain R5255 and OCN473 was used for transformant sequencing.

The R5362 strain (*comC0, SpcomEC*$^{R261A\text{-}alfa349}$), was generated as described for strain R5355, except that primer pairs OCN484-MD82 and MD83-OMB12 were used for initial PCRs on strain R5255.

The R5363 strain (*comC0, hexA::spc, SpcomEC*$^{R261A}$) was generated by transforming R5281 with gDNA from R1843[5] and selecting for spectinomycin resistance.

To generate the R5366 strain (*comC0, SpcomEC*$^{D457K}$*, comFC*$^{F16D}$), R5283 was transformed with a DNA fragment containing the *comFC*$^{F16D}$ mutation (generated as previously described) without selection and positive clones were determined by PCR amplification with the CJ359-CJ364 primer pair and sequencing with the CJ362 primer.

The R5367 strain (*comC0, comFA*$^{W273D}$) was generated as described for strain R5064, except that primer pairs CJ868-MD102 and MD103-CJ873 were used for initial PCRs, and sequencing was carried out with CJ344 primer.

To generate the R5368 strain (*comC0, hexA::spc, comFA*$^{W273D}$*, SpcomEC*$^{R261A}$), R5363 was transformed with a DNA fragment containing the *comFA*$^{W273D}$ mutation (generated as previously described) without selection and positive clones were determined by PCR amplification with the CJ359-CJ364 primer pair and sequencing with the CJ344 primer.

The R5372 strain (*comC0, comFC*$^{F16D\text{-}K83D}$) was generated as described for strain R5064, except that primer pairs CJ868-MD80 and MD81-CJ873 were used for initial PCRs on strain R5191, and sequencing was carried out with CJ364 primer.

To generate the R5393 strain (*comC0, comEC*$^{R261A}$*, comFC*$^{F16D}$), R5281 was transformed with a DNA fragment containing the *comFC*$^{F16D}$ mutation (generated as previously described) without selection and positive clones were determined by PCR amplification with the CJ359-CJ364 primer pair and sequencing with the CJ362 primer.

To generate the R5395 strain (*comC0, comFA*$^{Y59D+W273A}$), R5278 was transformed with a DNA fragment containing the *comFA*$^{Y59D}$ mutation (generated as previously described) without selection and positive clones were determined by PCR amplification with the CJ359-CJ364 primer pair and sequencing with the CJ359 primer.

To generate the R5431 strain (*comC0, comEC*$^{D457K\text{-}alfa349}$*, comFC*$^{F16D}$), R5361 was transformed with a DNA fragment containing the *comFC*$^{F16D}$ mutation (generated as previously described) without selection and positive clones were determined by PCR amplification with the CJ359-CJ364 primer pair and sequencing with the CJ362 primer.

To generate the R5432 strain (*comC0, comEC*$^{R261A\text{-}alfa349}$*, comFC*$^{F16D}$) and the R5433 strain (comC0, comEC$^{R261A\text{-}alfa349}$, comFA$^{W273D}$), R5362 was transformed with a DNA fragment containing respectively the *comFC*$^{F16D}$ mutation (generated as previously described) or the *comFA*$^{W273D}$ mutation (generated as previously described) without selection and positive clones were determined by PCR amplification with the CJ359-CJ364 primer pair and sequencing with the CJ362 primer.

**Construction of *H. pylori* mutant strains.**

All plasmid constructions were carried out using a pJET2.1 backbone and sequence- and ligation-independent cloning[10]. For the ectopic expression of *HpcomEC-FLAG* fusions driven by the *HpcomEC* promoter, the region harbouring the *HpcomEC* gene and its promoter was amplified from *H. pylori* 26695 genomic DNA (gDNA) using a downstream primer coding for the tag and cloned between a 200 bp sequence upstream of the *rdxA* locus and a non-polar chloramphenicol resistance cassette followed by the downstream 200 bp flanking the *rdxA* locus. For the ectopic expression of *HpcomEC-2alfa* from the strong *ureA* promoter, the *HpcomEC* open reading was amplified from the genomic DNA (gDNA) of *H. pylori* 26695 using a downstream primer coding for two alfa tags[9] and cloned into a plasmid between a 200 bp sequence upstream of the *ureA* ORF and a non-polar chloramphenicol resistance cassette followed by the downstream 200 bp flanking the *ureA* gene.

To delete the *HpcomEC* locus, a non-polar kanamycin resistance cassette was amplified by PCR and inserted into the plasmid pJET2.1 using SLIC, flanked by 200 bp upstream and downstream of the *HpcomEC* gene. The resulting constructs were then introduced into *H. pylori* through natural transformation as described below. In all cases at least two independent clones were selected for each construct. For a detailed list of all strains, primers and plasmids used to generate mutant constructs, see Table S3.

### Supplementary results and discussion

#### Altered expression of *spcomFC* in split-luc strains

Western blot analysis of *spcomFA-lgbit smbit-spcomFC* strains revealed that the expression of *smbit-spcomFC* was altered in these strains (Figure S6E). In addition, these strains displayed lower transformation efficiency than isogenic parental strains (Figure S6F). In a wildtype context, the ribosome-binding site (RBS) of *spcomFC* is present in the sequence of the *spcomFA* open reading frame. To construct these strains, we duplicated the *spcomFC* RBS downstream of the *lgbit* sequence associated with *spcomFA,* essentially transcriptionally decoupling the *spcomFA* and *spcomFC* transcripts. It may be that this decoupling results in a lower expression of *spcomFC* in these strains, and resulting impact on transformation efficiency. The reduction in transformation efficiency is greatest in the mutant strains (Figure 6F), which may reflect that tagging these proteins has a minor impact on the integrity of the tripartite complex, which is accentuated in mutant conditions. However, despite displaying a lower expression of *spcomFC*, and a significant decrease in transformation efficiency, the *Sp*ComFC$^{K83D}$ mutant, which is not proposed to impact *Sp*ComFA-FC interaction, shows maintained luminescence in the split-luc assay (Figure 4E). This is in contrast with the *Sp*ComFA$^{Y59D}$ and *Sp*ComFC$^{F95D}$ mutants, and serves to validate the pertinence of the loss of luminescence in these mutants.

#### Altered expression of *spcomFC* in *comFA*$^{stop}$ strain

Western blot analysis of the *spcomFA*$^{stop}$ strain revealed that the expression of *spcomFC* was also reduced in this strain (Figure S7A). As a result, it was not possible to rule out that the reduction in transformation efficiency observed in a *spcomFA*$^{stop}$ mutant was due to a reduction in *spcomFC* expression rather than the loss of *spcomFA*. To address this, we complemented the *spcomFA*$^{stop}$ mutant with ectopic expression of *spcomFC* from the *CEPlac* platform[11] and compared expression of *spcomFA* and *spcomFC* and transformation efficiency. Western blot analysis confirmed the complementation of *spcomFC* expression in this strain (Figure S7D), and transformation assays revealed that complementing *spcomFC* expression in the *spcomFA*$^{stop}$ mutant did not increase transformation efficiency (Figure S7E). This confirmed that the loss of *Sp*ComFA itself resulted in the observed decrease in transformation efficiency, validating that both *Sp*ComFA and *Sp*ComFC are crucial for pneumococcal transformation.

### Supplementary Figures

**Figure S1: Illustration of the ssDNA interaction surface on the tripartite *Sp*ComEC/FA/FC model.** (A) Charge representation of *Sp*ComEC interaction zones with *Sp*ComFA/FC, shown from below. (B) Conservation of *Sp*ComEC interaction zone in 10 firmicute species (Table S1), shown from below. (C) Charge representation of *Sp*ComFA interaction zones with *Sp*ComEC, *Sp*ComFC and ssDNA. (D) Conservation of *Sp*ComEC interaction zone in 10 firmicute species (Table S1). (E) Charge representation of *Sp*ComFC interaction zones with *Sp*ComEC, *Sp*ComFA and ssDNA. (F) Conservation of *Sp*ComEC interaction zone in 10 firmicute species (Table S1).

**Figure S2: Conservation of tripartite ComEC/FA/FC model in firmicutes.** Structural models of tripartite transformation membrane transport complex composed of ComEC (purple), ComFA (green) and ComFC (red) interacting with ssDNA (black) in nine other firmicute species. EC, ComEC, FA, ComFA, FC, ComFC. UniProtKB accession numbers of proteins used and model details can be found in Table S2. Polarity of ssDNA in the model is shown. (A) *Bacillus subtilis*. (B) *Lactococcus lactis.* (C) *Lactococcus sakei.* (D) *Staphylococcus aureus.* (E) *Streptococcus epidermidis.* (F) *Streptococcus mutans.* (G) *Streptococcus salivarius.* (H) *Streptococcus sanguinis.* (I) *Streptococcus thermophilus.*

**Figure S3: Validation of expression of *Sp*ComEC mutants in *S. pneumoniae.*** (A) Structural model of *Sp*ComEC protein highlighting domains and cytoplasmic loop into which alfa tag is inserted, after AA349. (B) Transformation efficiency of *Sp*ComEC$^{alfa349}$ strain compared to wildtype in saturating and limiting tDNA concentrations. tDNA identity and concentrations, as well as representations and statistical analyses as in Figure 2FG. (C) Time-course Western blot of *Sp*ComEC$^{alfa349}$ strain (R5255) after competence induction through CSP addition at t = 0 min. *Sp*ComEC$^{alfa349}$ detected using anti-alfa antibodies, competence specific SsbB and constitutive SsbA detected using anti-SsbB antibodies as a control. (D) Time-course Western blot of *Sp*ComEC$^{alfa349}$ strain as in panel C but over a shorter time window. (E) Evolutionary conservation of *Sp*ComEC residues derived from a multiple sequence alignment (see Methods), mapped from white (most variable) to red (most conserved) Positions which were experimentally mutated are shown in sticks. (F) Validation of pneumococcal *Sp*ComEC mutant expression in *Sp*ComEC$^{alfa349}$ strain by Western blot. Samples taken 8 min after competence induction by CSP addition. *Sp*ComEC$^{alfa349}$ detected using anti-alfa antibodies, SsbB detected using anti-SsbB antibodies as a control. Strains used: *SpcomEC*, R1501; wt, R5255; Y116E-Y157E, R5355; F152D, R5356; R292M, R5357; R292K, R5358; K297E, R5359.

**Figure S4: Schematic representations of pneumococcal transformation conditions in saturating and limiting tDNA concentrations.** (A) Transformation of competent pneumococcal cells in saturating concentrations of transforming ssDNA. In this case, it is assumed that all competent cells internalize multiple ssDNA fragments through the *Sp*ComEC transformation channel, resulting in integration of the *rpsL41* selectable resistance marker being integrated into the chromosome of the vast majority of cells. From this, results a transformation efficiency of between 50-100%, since resolution of a singly transformed chromosome will generate one sensitive and one resistant daughter cell. This condition ensures maximal transformation efficiency, but can mask the minor impacts of specific mutations on transformation efficiency. (B) In contrast, when the concentration of transforming ssDNA is limiting (100-fold less than saturating), the decrease in tDNA in the medium will result in only a fraction of cells internalizing ssDNA via *Sp*ComEC, and in most cases a single ssDNA fragment will be internalized. This results in a lower transformation efficiency, and may reveal subtle effects of specific mutations on transformation efficiency.

**Figure S5: Validation of production and stability of *Hp*ComEC mutants in *H. pylori.*** (A) Structural alignment of *Sp*ComEC (blue) and *Hp*ComEC, coloured by evolutionary conservation derived from a multiple sequence alignment (see Methods), from white (most variable) to red (most conserved). Corresponding positions are shown in sticks: *Sp*ComEC$^{Y116}$ and *Hp*ComEC$^{Y82}$ (left panel), *Sp*ComEC$^{R292}$ and *Hp*ComEC$^{R239}$ (right panel); ssDNA is shown in black. (B) *Hp*ComEC structural model, coloured by evolutionary conservation derived from a multiple sequence alignment (see Methods), from white (most variable) to red (most conserved). The Y60 position which was experimentally mutated is shown in sticks. ssDNA is shown in black. (C) Validation of *Hp*ComEC mutants in *Hp*ComEC-FLAG over-expressed from $P_{ureA}$ promoter. *Hp*ComEC-FLAG detected using anti-FLAG antibodies, membrane-associated protein MotB detected using anti-MotB antibodies as a control. Strains used: $P_{ureA}$-*HpcomEC-alfa*, 1435; wt, 26695; R239M, 1439; R239K, 1454; Y60E-Y82E, 1456.

**Figure S6: Validation of production and stability of *Sp*ComFA and *Sp*ComFC mutants in *S. pneumoniae*.** (A) Time-course Western blot of wt strain (R1501) after competence induction through CSP addition at t = 0 min. *Sp*ComFA detected using anti-*Sp*ComFA antibodies, *Sp*ComFC detected using anti-*Sp*ComFC antibodies, competence specific SsbB and constitutive SsbA detected using anti-SsbB antibodies as a control. (B) Validation of *Sp*ComFC mutant production by Western blot. Samples taken 10 min after competence induction by CSP addition. *Sp*ComFC detected using anti-*Sp*ComFC antibodies, SsbB detected using anti-SsbB antibodies as a control. Strains used: *wt*, R1501; Δ*FC*, R5099; F95D, R5153; K83D, R5191; R156D, R5238; F16D, R5279; Δ12-26, R5280. (C) Validation of *Sp*ComFA mutant production by Western blot. Samples taken 10 min after competence induction by CSP addition. *Sp*ComFA detected using anti-*Sp*ComFA antibodies, SsbB detected using anti-SsbB antibodies as a control. Strains used: *wt*, R1501; Δ*FA*, R5114; Y59D, R5064; W273A, R5278; W273D, R5367. (D) Validation of *Sp*ComEC mutant production in *Sp*ComEC$^{alfa349}$ strain by Western blot. Samples taken 8 min after competence induction by CSP addition. *Sp*ComEC$^{alfa349}$ detected using anti-alfa antibodies, SsbB detected using anti-SsbB antibodies as a control. Strains used: *SpcomEC*, R1501; wt, R5255; E267K, R5282; R261A, R5281; D457K, R5283. (E) Validation of protein production of split luciferase strains by Western blot. Samples taken 8 min after competence induction by CSP addition. *Sp*ComFA detected using anti-*Sp*ComFA antibodies, *Sp*ComFC detected using anti-*Sp*ComFC antibodies, SsbB detected using anti-SsbB antibodies as a control. Strains used: *wt*, R1501; Δ*FC*, R5099; Δ*FA*, R5114; *spcomFA-lgbit*, R5194; *spcomFA-lgbit smbit-comFC*, R5162; *spcomFA-lgbit smbit-comFC* Δ*EC*, R5252; *spcomFA*$^{Y59D}$*-lgbit smbit-comFC*, R5179; *spcomFA-lgbit smbit-comFC*$^{F95D}$, R5240; *spcomFA-lgbit smbit-comFC*$^{K83D}$, R5257. (F) Transformation efficiencies of split-luc strains with saturating tDNA. Strains as in panel E with R5252 removed since *spcomEC*$^{-}$ mutants do not transform.

**Figure S7: Validation of production and stability of pneumococcal proteins in double mutants.** (A) Validation of *Sp*ComFA and *Sp*ComFC protein production in double mutants by Western blot. Samples taken 8 min after competence induction by CSP addition. *Sp*ComFA detected using anti-*Sp*ComFA antibodies, *Sp*ComFC detected using anti-*Sp*ComFC antibodies, SsbB detected using anti-SsbB antibodies as a control. Strains used: *wt*, R1501; Δ*FC*, R5099; Δ*FA*, R5114; *spcomFA*$^{Y59D}$ *spcomFC*$^{K83D}$, R5251; *spcomFC*$^{K83D+R156D}$, R5270; *spcomFA*$^{Y59D}$ *spcomFC*$^{F95D}$, R5305; *spcomFA*$^{W273A}$ *spcomFC*$^{F16D}$, R5307; *spcomFA*$^{Y59D}$ *spcomFC*$^{F16D}$, R5309; *spcomFC*$^{F16D+K83D}$, R5372; *spcomFA*$^{Y59D+W273A}$, R5395. This Western blot revealed that *Sp*ComFC is produced at lower levels in a Δ*spcomFA* strain, which is further explored and discussed in Figure S7DE and Supplementary Information. (B) Validation of *Sp*ComFA and *Sp*ComEC$^{alfa349}$ protein production in double mutants by Western blot. Samples taken 8 min after competence induction by CSP addition. *Sp*ComFA detected using anti-*Sp*ComFA antibodies, *Sp*ComEC$^{alfa349}$ detected using anti-alfa antibodies, SsbB detected using anti-SsbB antibodies as a control. Strains used: *wt*, R1501; Δ*FA*, R5114; *spcomFA*$^{W273D}$ *spcomEC*$^{alfa349-R261A}$, R5433. (C) Validation of *Sp*ComFC and *Sp*ComEC$^{alfa349}$ protein production in double mutants by Western blot. Samples taken 8 min after competence induction by CSP addition. *Sp*ComFC detected using anti-*Sp*ComFC antibodies, *Sp*ComEC$^{alfa349}$ detected using anti-alfa antibodies, SsbB detected using anti-SsbB antibodies as a control. Strains used: *wt*, R1501; Δ*FC*, R5099; *spcomFC*$^{F16D}$ *spcomEC*$^{alfa349-D457K}$, R5431; *spcomFC*$^{F16D}$ *spcomEC*$^{alfa349-R261A}$, R5432. (D) Complementation of *spcomFC* expression in a *comFA*$^{stop}$ genetic context, revealed by Western blot. *Sp*ComFA detected using anti-*Sp*ComFA antibodies, *Sp*ComFC detected using anti-*Sp*ComFC antibodies, SsbB detected using anti-SsbB antibodies as a control. Strains used: *wt*, R1501; Δ*FA*, R5114; Δ*FA* *CEP*$_{lac}$-*comFC*, R5121. (E) Transformation efficiency of *spcomFA*$^{stop}$ mutant and *spcomFC* complemented strain in presence of IPTG. Strains used as in panel D, with saturating tDNA concentrations.

**Figure S8: Dual mutations targeting different protein-protein and protein-ssDNA interactions with *Sp*ComFC have cumulative effects on transformation.** (A) Schematic representation of tripartite model of tDNA membrane transport, showing key *Sp*ComFC interaction zones with ssDNA and partner proteins. (B) Highlight of tripartite structural model of tDNA membrane transport, with three residues shown, one key to each interaction of *Sp*ComFC tested for this model. (C) Transformation efficiency of single or double *Sp*ComFC interaction mutants when transformed with saturating concentrations of tDNA. tDNA identity and concentration as in Figure 2F. Strains used: *wt*, R1501; $FA^{Y59D}$, R5064, $FC^{K83D}$, R5191; $FC^{F16D}$, R5279; $FA^{Y59D}$ $FC^{K83D}$, R5251; $FA^{Y59D}$ $FC^{F16D}$, R5309. Error bars representative of triplicate repeats, with individual data points shown. Student's t-tests were used to compare transformation efficiencies of mutant strains with wildtype, unless shown by horizontal line. (D) Transformation efficiency of single and double *Sp*ComFC interaction mutants when transformed with limiting concentrations of tDNA. tDNA identity and concentration as in Figure 2G. Data representation as in panel C.

**Table S1:** Conservation of ComEC, ComFA and ComFC in a selection of strains chosen to represent the diversity of transformable species across the bacterial kingdom[12]. Codes refer to UniProtKB accession numbers of proteins for each species. The presence of the predicted ComFC N-terminal 'hook' structure is also shown.

**Table S2:** Information about AlphaFold3 models of ComEC/FA/FC with ssDNA in ten firmicute species, including *S. pneumoniae*. For each species, the table shows UniProt accession numbers of the three proteins, AlphaFold3 model confidence scores (ipTM and pTM), RMSD measuring the deviation between the AlphaFold3 model in this species and the *S. pneumoniae* model, interacting surface areas (in Å²) for each pairwise protein-protein interface in the model and sequence identity with pneumococcal sequences.

**Table S3**: Strains and primers used in this study.
